# Transposon accumulation lines uncover histone H2A.Z-driven integration bias towards environmentally responsive genes

**DOI:** 10.1101/447870

**Authors:** Leandro Quadrana, Mathilde Etcheverry, Arthur Gilly, Erwann Caillieux, Mohammed-Amin Madoui, Julie Guy, Amanda Bortolini Silveira, Stefan Engelen, Victoire Baillet, Patrick Wincke, Jean-Marc Aury, Vincent Colot

**Affiliations:** Institut de Biologie de l’Ecole Normale Supérieure (IBENS), Centre National de la Recherche Scientifique (CNRS), Institut National de la Santé et de la Recherche Médicale (INSERM), Ecole Normale Supérieure, PSL Research University, Paris, France.; Genoscope, Institut de biologie François-Jacob, Commissariat à l’Energie Atomique (CEA), Université Paris-Saclay, F-91057 Evry, France.; Génomique Métabolique, Genoscope, Institut de biologie François Jacob, CEA, CNRS, Univ Evry, Université Paris-Saclay, 91057 Evry, France.; Human Genetics, Wellcome Trust Sanger Institute, Hinxton, UK.; Translational Research Department, Institut Curie, PSL Research University, Paris, France.

## Abstract

Inherited transposition events are important drivers of genome evolution but because transposable element (TE) mobilization is usually rare, its impact on the creation of genetic variation remains poorly characterized. Here, we used a population of *A. thaliana* epigenetic recombinant inbred lines (epiRILs) to characterize >8000 de novo insertions produced by several TEs families also active in nature. Integration was strongly biased towards genes, with evident deleterious effects. Biases were TE family-specific and associated with distinct chromatin features. Notably, we demonstrate that the histone variant H2A.Z guides the preferential integration of *Ty1/Copia* LTR-retrotransposons within environmentally responsive genes and that this guiding function is evolutionary conserved. Finally, we uncover an important role for epigenetic silencing in exacerbating or alleviating the effects of TE insertions on target genes. These findings establish chromatin as a major determinant of the spectrum and functional impact of TE-generated mutations, with important implications for adaptation and evolution.

## INTRODUCTION

Transposable elements (TEs) are sequences that self-propagate and accumulate to various levels in the genome of eukaryotic species. TEs fall into two broad classes, depending on their mechanism of transposition: DNA transposons, which move through a “cut and paste” mechanism and Long Terminal Repeat (LTR) as well as non-LTR retrotransposons, which move through an RNA intermediate that is reverse transcribed. The content of the different classes and subclasses of TEs varies markedly across species. Thus, mammalian genomes are laden with non-LTR retrotransposons whereas plant genomes are populated by LTR retrotransposons and DNA transposons mainly (Lisch 2013; Chuong, Elde, and Feschotte 2016). However, most of these sequences are relics of once active TEs and although TEs are arguably among the main drivers of the evolution of genome size, organization and function (Chuong et al., 2016), we still know little about their contribution to the generation of somatic or heritable mutations in extant genomes (Huang, Burns, and Boeke 2012; Sultana et al. 2017). This situation stems in large part from the fact that transposition is typically rare in nature, notably because of the strong epigenetic silencing mechanisms, such as DNA methylation in plants and mammals, that target TEs (Slotkin and Martienssen 2007).

With the advent of population genomics, it is now possible to make use of natural variation in the number and distribution of recent TE insertions to gain information about the set of TEs likely mobile in extant genomes. However, because of the deleterious effects typically associated with TE mobilization, natural selection prevents the proper assessment of the complete range of the mutations generated by transposition (Huang, Burns, and Boeke 2012; Sultana et al. 2017; Quadrana et al. 2016).

Here, we have exploited a population of *A. thaliana* epigenetic Recombinant Inbred Lines (epiRILs; Johannes et al., 2009) to obtain a first comprehensive assessment of the rate, spectrum, properties and genome-wide distribution of heritable mutations generated by TEs. The epiRILs were derived from a cross between a wild type and a near isogenic parent in which transposition was kick-started for several TEs as a result of compromised DNA methylation (Johannes et al., 2009; Figure 1A). Based on whole genome sequencing for more than 100 epiRILs and the characterization of >8000 *de novo* heritable insertions, we first establish that the epirRILs are operationally similar to mutation accumulation (MA) lines (Denver et al. 2009; Keightley et al. 2009; Zhu et al. 2014). We then show that two DNA transposons and one LTR-retrotransposon that are among the most active in nature target non-overlapping sets of genes and in relation to distinct chromatin features. We further demonstrate that the preferential targeting of environmentally responsive genes by LTR-retrotransposons of the *Ty1/Copia* superfamily is guided by the histone variant H2A.Z. We also provide evidence that this guiding function is ancestral, as it is conserved from plants to yeast and propose a plausible mechanistic explanation for the invasive success of *Ty1/Copia* retrotransposons exclusively within plant genomes. Finally, we show that epigenetic silencing of new insertions soon after their occurrence can either alleviate or exacerbate their effects on gene transcription. Our findings illustrate the role of chromatin as a major determinant of the spectrum and impact of TE-generated mutations, with important implications for adaptation as well as the evolutionary success of TEs and the organisms in which they reside.

**Figure 1.**
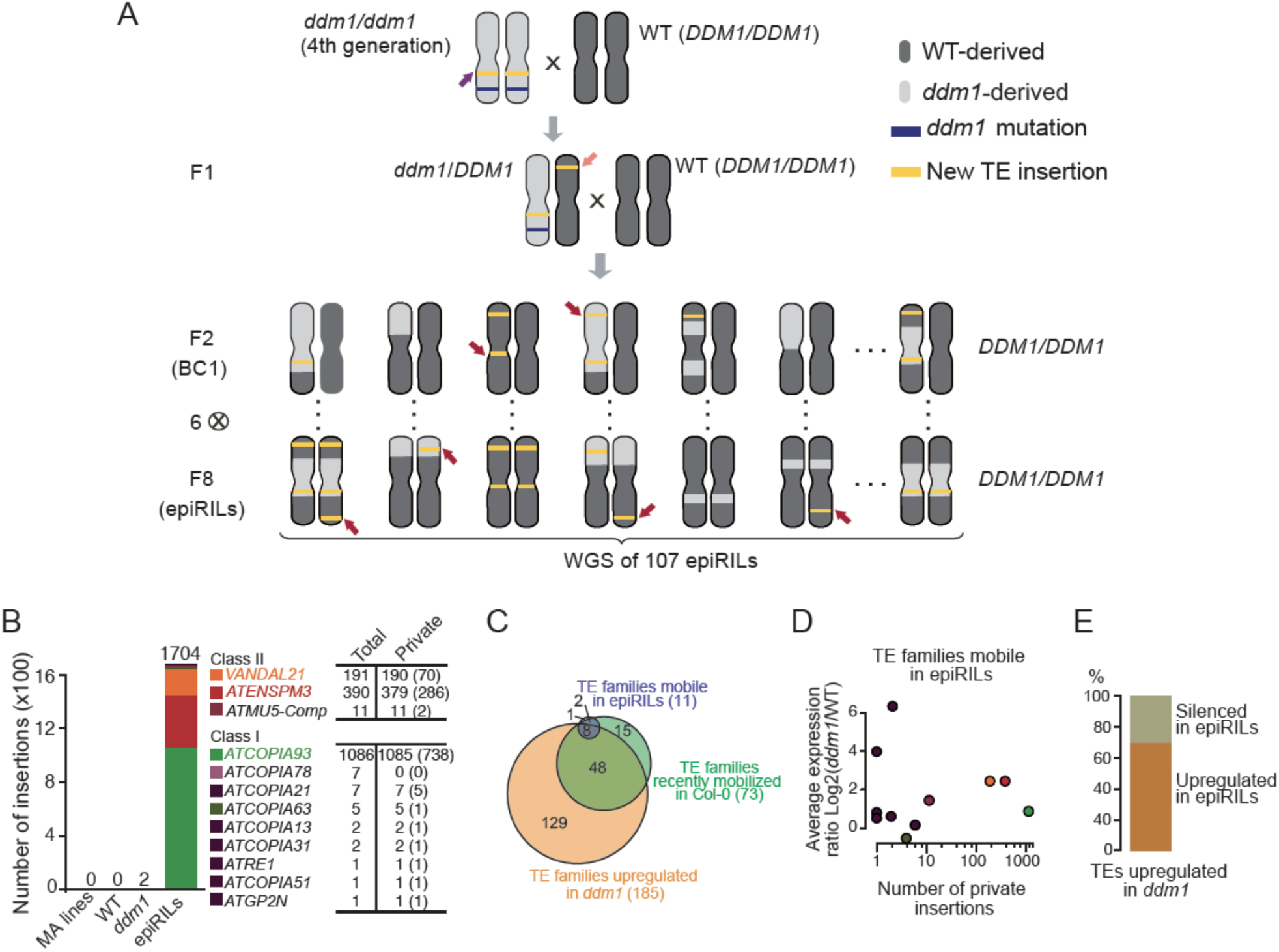
TE insertions accumulate in the epiRILs. **A.** Crossing scheme used to generate the epiRIL population. **B.** Number and identity of insertions accumulated in wild-type, *ddm1* and the epiRILs. Number of heterozygous insertions are indicated within brackets. **C.** Overlap between TE families transcriptionally reactivated in *ddm1*, those potentially mobile in Col-0 and those that transposed in at least one epiRIL. **D.** Relation between number of private TE insertions and average expression ratio (*ddm1*/wild-type) for the 11 TE families that transposed in the epiRILs. **E.** Proportion of reference TE sequences upregulated in *ddm1* and either silenced or stably upregulated in the epiRILs.

## RESULTS

### TE insertions accumulate in the epiRILs

We produced high quality whole genome sequencing data for 107 epiRILs at generation F8 as well as of close relatives of the wild type and *ddm1* parental lines (Figure 1A) using Illumina mate-pair libraries built from size-selected genomic fragments of ∼5.5 kb. Mate-pair reads were mapped to the reference genome sequence (accession Col-0) and non-reference (i.e. *de novo)* insertions were detected based on mapping discordance between mate-pair reads and using also split reads (Gilly et al., 2014; Quadrana et al., 2016; Figure S1A). Thanks to the large physical distance between mate-pair reads, the complete sequence of the *de novo* insertions was also determined, thus enabling the identification of the exact or most likely donor TE in each case. Consistent with the low frequency of TE mobilization in nature (Quadrana et al. 2016) there were no *de novo* TE insertions in the two wild type siblings sequenced nor in five *A. thaliana* MA lines (Ossowski et al. 2010). In contrast, two non-reference TE insertions were detected in the sequenced *ddm1* individual and many more in the epiRILs. Specifically, 95% of the 107 epiRILs harbored between 1 and 97 heritable *de novo* insertions and only 8 had none (Figure S1B). Almost all (98.7%) of these insertions were private and therefore did not occur in the *ddm1* parent but rather either in the inflorescences of the F1 individual used for the backcrossing or during the propagation of the epiRILs, starting from the F2 (Figure 1A). In fact, 1107 out of the 1670 private insertions detected in total were heterozygous (Figure 1B, S1B), which is strong evidence that transposition was ongoing in the epiRILs. Given that they were propagated by repeated selfing and single seed descent, the epiRILs could therefore be defined as TE accumulation (TEA) lines. In other words, notwithstanding the possibility of excision in the case of DNA transposons (see below), TE insertions that occurred during the propagation of the epiRILs and that were potentially heritable should be lost or fixed in a neutral fashion through random segregation, except for those disrupting essential genes.

Overall, we detected private insertions for two DNA transposon and eight retrotransposon families as well as for three composite DNA transposons (Figure 1B). These composite elements were uncovered thanks to our mate-pair sequencing approach and are made up for two of them of a protein-coding gene of unknown function flanked by two distinct truncated copies of *ATENSPM3*. The third composite DNA transposon is more complex and contains several truncated TEs, the largest of which is related to the *ATMU5* family (Figure S1C). The number of transposition events varied considerably among the eleven TE families, with the two DNA transposons *VANDAL21* and *ATENSPM3* (together with the two composite *ATENSPM3* elements) and the LTR-containing retrotransposon *ATCOPIA93* contributing 11.2%, 22.5% and 64.4% of the *de novo* insertions, respectively (Figure 1B).

We showed previously that 73 of the 326 TE families annotated in the *A. thaliana* genome transposed in the recent past in the reference accession Col-0 (Quadrana et al. 2016), which was used to derive the epiRIL population. Consistent with this finding the 10 annotated TE families with private insertions in the epiRILs are also mobile in nature and belong to the Col-0 mobilome. Nonetheless, the epiRIL mobilome is only a small subset of that of Col-0, which indicates that the widespread loss of DNA methylation induced by *ddm1* does not translate in an equally widespread remobilization of TE families. To determine if differences in the level of ddm1-induced transcriptional reactivation between TE families could explain at least in part the limited mobilome of the epiRILs, we performed RNA-seq on whole seedlings for five epiRILs as well as one sibling of the two parents. Between 731 and 1056 (considering unique or multiple mapping reads, respectively) TEs, mostly full-length, were significantly overexpressed at least 2-fold (*P*<0.05) in *ddm1* compared to wild type, consistent with previous RNA seq data sets (Figure S2). Moreover, >75% of the 73 TE families that compose the Col-0 mobilome were upregulated in *ddm1,* including eight of the 11 TE families that are part of the epiRIL mobilome (Figure 1C). However, levels of upregulation in *ddm1* did not correlate with the number of private insertions in the epiRILs (Figure 1D). Finally, 72% of the ddm1-upregulated TEs for which expression could be ascertained unambiguously in the epiRILs remained transcriptionally active when derived from the *ddm1* parent (Figure 1E). Thus, levels of transcription in seedlings did not reflect transposition activity. Whether this also holds true in the cells, starting from the zygote, that will ultimately pass on their DNA to the next generation remains to be determined, as these cells are where transposition must occur for insertions to be inherited.

### Invasion dynamics differ between TE families

The large number of private insertions for *VANDAL21*, *ATENSPM3* and *ATCOPIA93* enabled us to investigate in depth for each of these three TE families their mode and tempo of mobilization across generations. Mate-pair reads were first used to determine the sequence of the private insertions. In the case of *VANDAL21* and *ATCOPIA93*, almost all (99%) were identical to a single one of the 11 and four full-length copies present in the reference genome, respectively. This single copy is the same that was reported previously to be mobile in *ddm1* or another DNA methylation mutant background (Fu et al. 2013; Mirouze et al. 2009). In contrast, the identity of the *ATENSPM3* private insertions was more diverse, with 58% matching a single one of the nine full-length copies and the remaining private insertions matching either one of the two composite copies also present in the reference genome (Figure 2A). Each of these two composite *ATENSPM3* copies contains a single gene of unknown function with no similarity to any known transposase gene, indicating that they were likely mobilized in *trans*, presumably using the transposase encoded by the mobile full-length *ATENSPM3* reference copy. Thus, for each of these three TE families, a single *ddm1*-derived reference element is at the origin of most transposition events that accumulated in the epiRILs.

**Figure 2.**
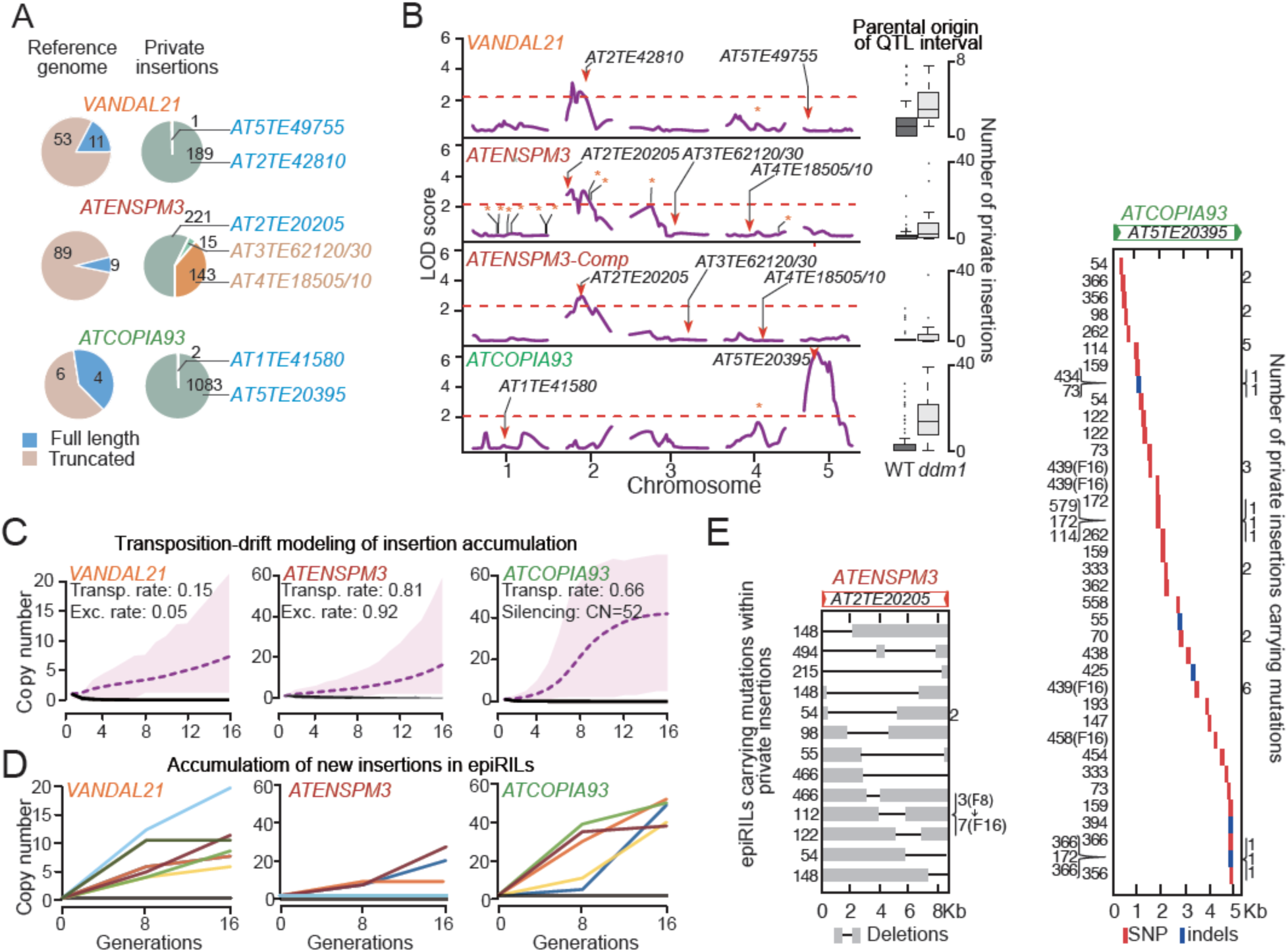
Invasion dynamics differ between TE families. **A.** Identification of donor copies for each transposition event. The number of truncated and full-length copies annotated in the *A. thaliana* reference genome is also shown. **B.** QTL mapping results for the mobilization of *VANDAL21, ATENSPM3*, the two composite *ATENSPM3* and *ATCOPIA93*. TEs annotated in the reference *A. thaliana* genome as well as donor and shared insertions are indicated by bars, arrowheads and stars, respectively. Box-plots on the right indicate for each TE family the numbers of insertions in the epiRILs in relation to the parental origin of the relevant QTL interval. **C.** Predicted dynamics of insertion accumulation for the three TE families based on the best fitted transposition-drift model. Estimates of the transposition and excision rates as well as of the number of TE insertions required for triggering concerted epigenetic silencing are indicated. **D.** Number of *VANDAL21, ATENSPM3*, and *ATCOPIA93* insertions observed at generation F8 and F16 in ten epiRILs. **E.** Mutations detected within private insertions of *ATENSPM3* and *ATCOPIA93*. Carrier epiRILs as well as the number of private insertions with the mutation are indicated in each case on the left and right side of each panel, respectively.

To confirm this conclusion and explore in more detail the genetic architecture underlying the large variation in the number of private insertions accumulated in the epiRILs, we performed a quantitative trait locus (QTL) analysis for each of the three TE families. We took advantage of the fact that the epiRILs are virtually isogenic but segregate hundreds of parental differentially methylated regions (DMRs) that can be used as *bona fide* genetic markers (Colomé-Taché et al, 2012; Cortijo et al., 2014) to identify loci whose epigenetic state is associated with transposition activity. Using the number of private insertions as the quantitative trait, this approach revealed a single QTL interval for each TE family, which included the full-length reference copy identified in the previous step (Figure 2B). Furthermore, the same QTL interval was obtained whether all *ATENSPM3* private insertions were considered or only those that match the two composite *ATENSPM3* present in the reference genome, thus demonstrating that these were mobilized in *trans* by the single full-length, *ddm1*-derived reference copy of *ATENSPM3*.

To determine if these observations were compatible with a transposition-drift scenario, we modeled TE mobilization for each of the three TE families based on three key parameters. These are the rate of transposition as well as of excision in the case of the DNA transposons, as this is an integral part of their mechanism of transposition, and copy-number dependent inhibition of transposition, as *ATCOPIA93* was shown to be epigenetically silenced when reaching around 40 copies (Marí-Ordóñez et al. 2013). For each TE family we selected the model that produced the best fit to the observed number and ratio of homozygous vs heterozygous private insertions at generation F8 and ran each model for another eight generations (F16; Figure S3A). The selected models predicted that in addition to the original reference TE donor, all new insertions are equally able to transpose, at a rate of 0.15, 0.81 and 0.66 new copy/per donor/per generation for *VANDAL21*, *ATENSPM3* and *ATCOPIA93*, respectively (Figure 2C and S3B). The best-fit models also predicted low and high rates of excision for *VANDAL21* and *ATENSPM3*, as was observed in maize for TEs belonging to the same two superfamilies *Mu* and *En/Spm*, respectively (Doseff, Martienssen, and Sundaresan 1990; Masson et al. 1987). In agreement with these predictions, the mobile reference copy of *VANDAL21* was almost always retained in the epiRILs, unlike that of *ATENSPM3*, which was systematically missing from the corresponding *ddm1*-derived chromosome interval (Figure S3C). Furthermore, indels compatible with *ATENSPM3* excision footprints were consistently detected across the genome and their number correlated positively with that of *ATENSPM3* private insertions (Figure S3D). Finally, our modeling predicted that transposition of *ATCOPIA93* should stop when reaching 52 copies (Figure 2C and S3B). To evaluate further the predictive power of our models, we sequenced ten epiRILs that were propagated for another 8 generations (F16). Consistent with numerical simulations (Figure 2C), *VANDAL21* and *ATENSPM3* continued to transpose in most of the lines with initial transposition activity, with no evidence of copy-number-dependent transposition inhibition. In contrast, although *ATCOPIA93* continued to transpose beyond F8, none of the F16 lines had accumulated more than 50 copies (Figure 2D). Furthermore, almost all copies were in the homozygous state at F16, consistent with the concerted epigenetic silencing expected at this copy number (Marí-Ordóñez et al. 2013).

Last, we took advantage of the fact that mobilized TEs tend to mutate during the transposition process to obtain direct evidence that in addition to the initial *ddm1*-derived donor copy, new insertions were also mobilized in the epiRILs. While all new insertions were identical to the mobile reference element for *VANDAL21*, approximately 5% of *ATENSMP3* and *ATCOPIA93* new insertions identified at F8 and/or F16 contained large internal deletions and point mutations or small indels, respectively (Figure 2E). Some mutations were carried by more than one new insertion within an epiRIL, but rarely between epiRILs. Notably, for one of the 10 epiRILs also sequenced at F16, we observed an increase in the number of *ATENSPM3* insertions containing the same mutation at that later generation (Figure 2E). These findings provided therefore further evidence that newly inserted copies were also mobilized in the epiRILs.

### TEs exhibit strong and diverse chromatin-associated insertion biases towards genes

As in nature (Quadrana et al. 2016), private insertions for *VANDAL21, ATENSPM3* and *COPIA93* were distributed evenly across the five chromosomes in the epiRILs, overall (Figure 3A). However, because the genomes of the epiRILs are epigenetic mosaics, each being composed on average of 25% *ddm1*- and 75% wild-type-derived segments (Colomé-Tatché et al, 2012), we searched for potential differences in insertion frequency associated with parental origin. For all three TE families, the percentage of insertions in *ddm1*-derived intervals was slightly higher than expected by chance at the whole genome level but much higher (35-55%) when considering the pericentromeric regions only (Figure 3A and S4A), which lose their heterochromatic features in *ddm1* as well as in subsequent generations (Soppe et al. 2002; Lippman et al. 2004; Colome-Tatche et al. 2012). These findings reinforced the notion that euchromatin is the preferred substrate for the integration of *VANDAL21, ATENSPM3* and *COPIA93*.

**Figure 3.**
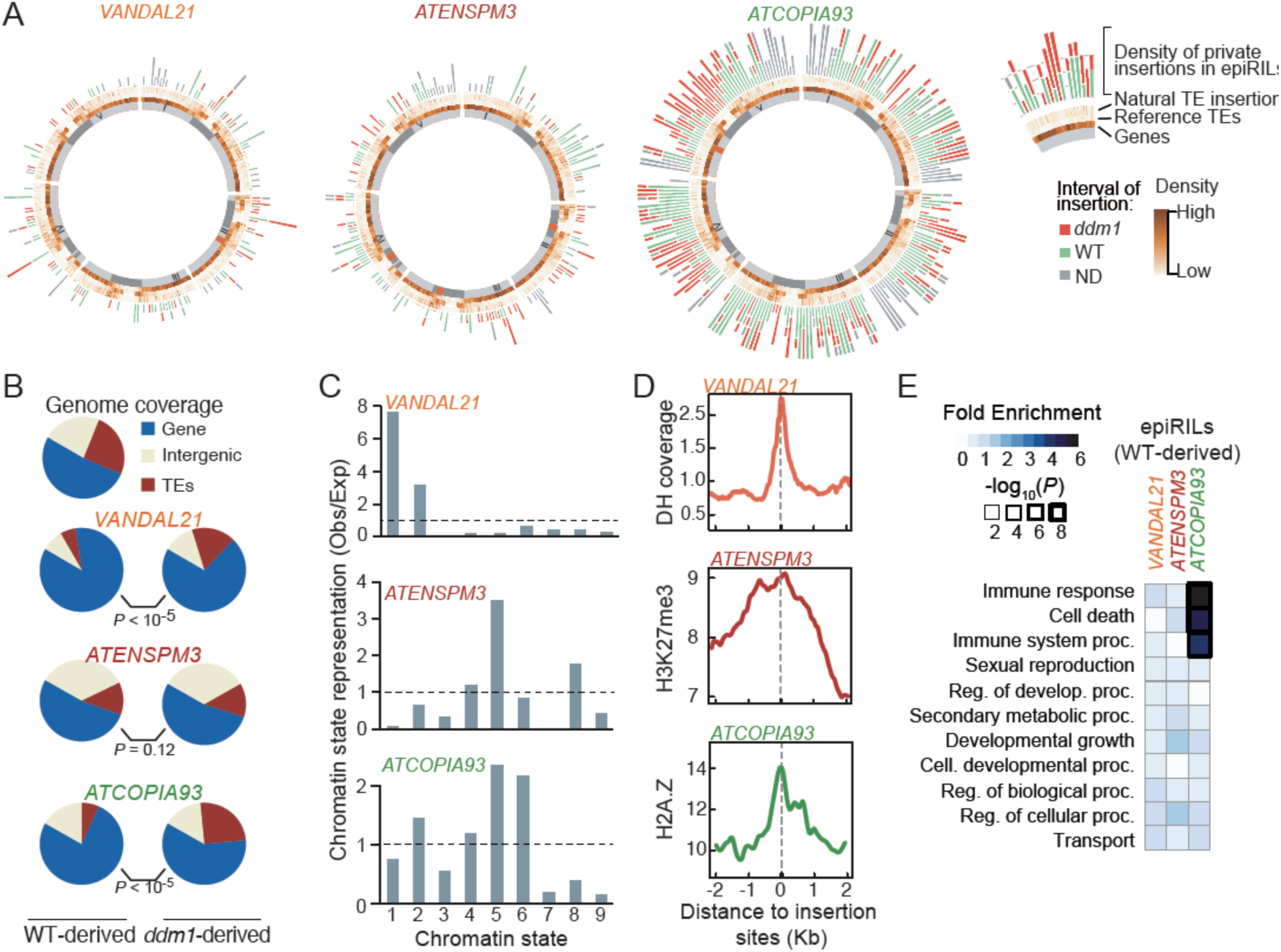
TEs exhibit strong and diverse chromatin-associated insertion biases towards genes. **A.** Circos representation of private TE insertions detected for *VANDAL21, ATENSPM3* and *ATCOPIA93*. The exterior circle represents the density of private insertions within wild-type- and *ddm1*-derived intervals. Density of non-reference (natural) TE insertions, reference TEs as well as genes are represented inwards in this order. **B.** Fraction of TE insertions in wild-type- and *ddm1*-derived intervals in genes, TEs and intergenic regions. Statistical significance for each comparison was obtained using the Chi-square test. **C.** Observed/expected ratio (Obs/Exp) of insertion sites within wild-type-derived regions in relation to the nine chromatin states defined in *A. thaliana*. Error bars represent the 95% confidence interval obtained by 1000 boots-traps. **D.** Coverage of DNAse hypersensitivity (DH; top panel), H3K7me3 (middle panel) and H2A.Z (bottom panel) around private insertion located within wild-type-derived regions for *VANDAL21*, *ATENSPM3* and *ATCOPIA93*, respectively. **E.** GO term analysis of genes with private TE insertions in the epiRILs.

Given that the methylome as well as the transcriptome of wild-type-derived intervals in the epiRILs are with few exceptions identical to their wild type parental counterparts (Colome-Tatche et al., 2012; Ito et al., 2015; Figure S2), we then used the private insertions located within these intervals to obtain information on integration preferences. Insertions were strongly overrepresented within or adjacent to genes for the three TE families (Figure 3B). However, while the fraction of essential genes (Lloyd et al. 2015) that were targeted was that expected by chance for *VANDAL21*, it was much lower for *ATENSPM3* and *ATCOPIA93*, suggesting that their integration, unlike that of *VANDAL21*, strongly affects the expression of target genes. Consistent with this interpretation, the fraction of essential genes targeted by *ATCOPIA93* was further reduced when only considering insertions in the homozygous state (Figure S4B). Unexpectedly though, we found an opposite pattern for *ATENSPM3* insertions within essential genes, which were less frequent in the heterozygous than the homozygous state. Given the high rate of excision associated with *ATENSPM3* mobilization, this last result suggested stronger deleterious effects after excision of *ATENSPM3*. Consistently, none of the 98 homozygous excision footprints detected in the epiRILs were located within essential genes (Figure S4B).

Using the nine main chromatin states defined in *A. thaliana* based on epigenomic maps obtained from diverse organs and tissues (Sequeira-Mendes et al. 2014), we found that *VANDAL21* mainly targeted promoters and 5’ UTRs of genes characterized as active (chromatin states 1&2; Figure 3C). In contrast, *ATENSPM3* and *ATCOPIA93* preferentially inserted within or close to genes that are typically marked by H3K27me3 and slightly enriched for the variant histone H2A.Z (chromatin state 5) and, in the case of *ATCOPIA93*, also in or adjacent to genes that were only enriched for that histone variant (chromatin state 6; Figure 3C).

Meta-analysis of insertion sites confirmed these findings and revealed that *VANDAL21* integration within wild-type-derived intervals tended to coincide with peaks of DNase I hypersensitivity (Figure 3D) at the transcriptional start site (TSS) of genes (Figure S4C). Furthermore, insertions were typically in the same orientation as the target genes (Figure S4C). These results support previous observations reported for *VANDAL21* in *ddm1* mutant plants (Fu et al.,2013) as well as in nature (Quadrana et al. 2016). In contrast, insertion sites were enriched for H3K27me3 in the case of *ATENSPM3* and for nucleosomal DNA as well as H2A.Z in the case of *ATCOPIA93* (Figure S4D). However, *ATCOPIA93* did not insert preferentially within well-positioned nucleosomes (Figure S4E; Lyons and Zilberman, 2017) nor within so-called “+1 nucleosomes”, which tend to incorporate H2A.Z (Henikoff and Smith 2015), but rather within genes (Figure S4F) that are enriched in *H2A.Z* throughout their body (Figure S4G). Such pattern of H2A.Z deposition is specific to plants and was shown previously to concern mainly responsive genes (Coleman-Derr and Zilberman 2012; Zahraeifard et al. 2018). In agreement, this class of genes was strongly overrepresented among *ATCOPIA93* targets (Figure 3E).

### H2A.Z directs the integration of *ATCOPIA93*

The *Ty1/copia* superfamily of LTR-retrotransposons is one of the main contributors of genome size inflation seen in many plant species (Huang, Burns, and Boeke 2012). We therefore explored further the function of chromatin in the integration of *ATCOPIA93* and first noted that the proportion of insertions within TE sequences was much higher in *ddm1*-derived than in wild-type-derived intervals (Figure 3B). This finding is entirely consistent with the fact that TE sequences tend to acquire H2A.Z when hypomethylated (Zilberman et al. 2008). Indeed, H2A.Z enrichment levels measured in the hypomethylated background *met1* that was used for this previous analysis were on average much higher, in *ddm1*-derived intervals, for TE sequences targeted by *ATCOPIA93* than for those that were not (Figure S4H).

To demonstrate the involvement of H2A.Z in the integration of *ATCOPIA93*, we crossed one epiRIL (epiRIL54, F8) containing 23 active (mainly heterozygous) *ATCOPIA93* copies to a wild type or an *hta9 hta11* double mutant parent, which lacks most H2A.Z (March-Díaz et al. 2008). Two F1 individuals were selfed in each case and two homozygous wild type as well as two homozygous double mutant F2 lines were selected (Figure 4A) to produce F3 progeny for TE-sequence capture (Quadrana et al. 2016). DNA was extracted from approximately 1000 F3 seedlings from each of the four F2 lines. Over 2000 new *ATCOPIA93* insertions were detected in each of the two wild type F3 progenies and less than twice that number in the two *hta9 hta11* F3 progenies (Figure 4B). In addition, while the strong *ATCOPIA93* integration preferences observed in the epiRILs was confirmed in wild type F3 seedlings, they were totally abolished in the *hta9 hta11* lines (Figure 4C, 4D, 4E). Moreover, the proportion of essential genes targeted by *ATCOPIA93* almost tripled in the double mutant (Figure 4F). These results demonstrated that H2A.Z acts at two levels, to promote *ATCOPIA93* retrotransposition and to guide integration within environmentally responsive genes. To investigate at the nucleosome-scale the guiding function of H2A.Z, we crossed the list of the *ATCOPIA93* insertion sites detected in the wild type F3 progeny with the list of well-positioned nucleosomes previously produced for *A. thaliana* (Lyons and Zilberman 2017). A major peak of integration was observed ∼55 bp away from the nucleosome dyad (Figure 4G). This position is a main point of contact between DNA and H2A or H2A.Z and it is also where these two histones differ by several amino acids (Suto et al. 2000; Zlatanova and Thakar 2008). These findings further support a direct role of H2A.Z in guiding *ATCOPIA93* integration.

**Figure 4.**
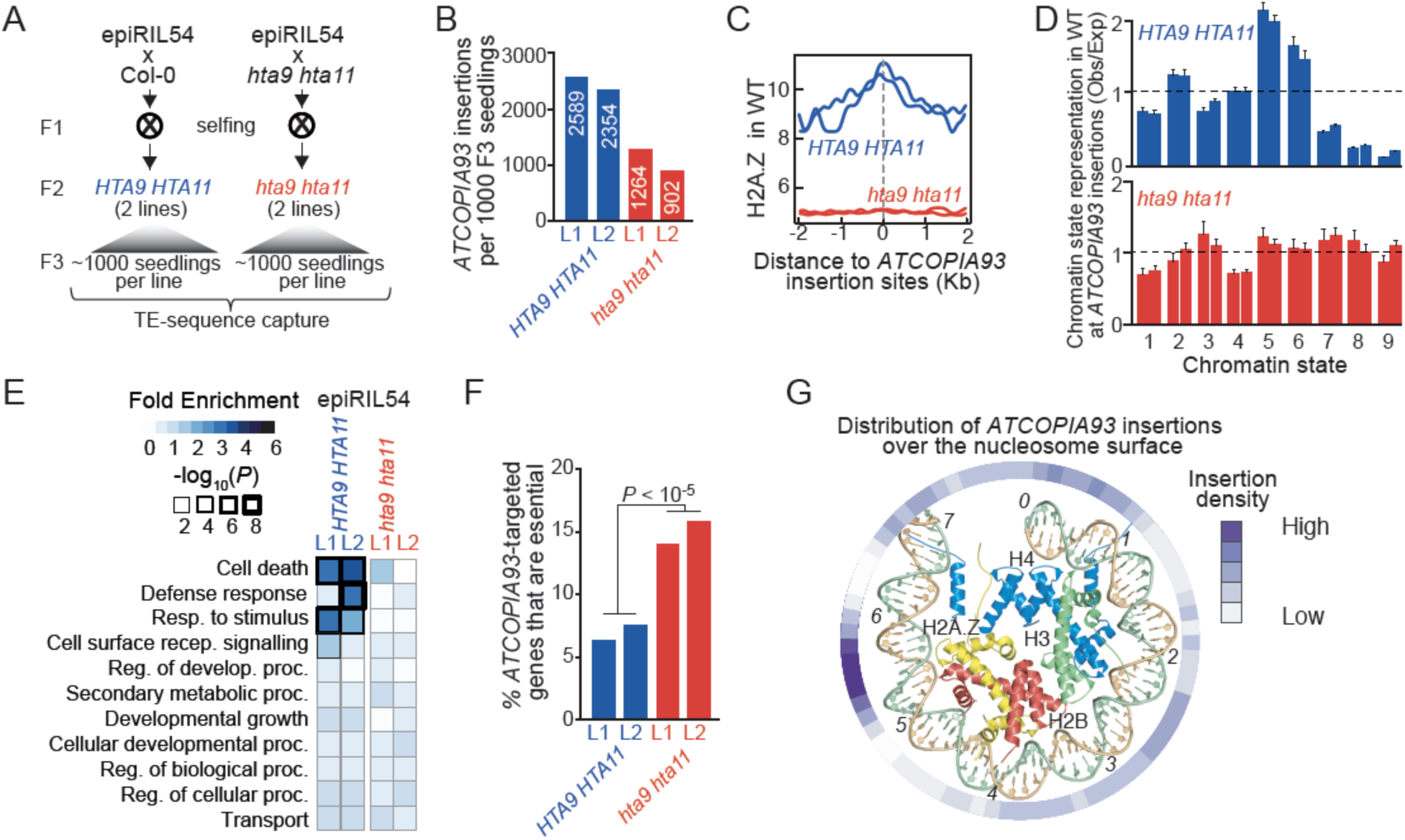
H2A.Z directs the integration of *ATCOPIA93*. **A.** Experimental strategy for determining the role of H2A.Z in the integration of *ATCOPIA93.* epiRIL54 was crossed with wild type or the double mutant *hta9 hat11.* Two wild type (*HTA9 HTA11* L1 and L2) and two double mutant (*hta9 hta11* L1 and L2) F2 plants were selected and 1000 F3 seedlings were collected in each case to perform TE-sequence capture. **B.** Number of new insertions detected in 1000 *HTA9 HTA11* and *hta9 hta11* F3 seedlings. **C.** Meta analysis of levels of H2A.Z around *ATCOPIA93* insertion sites. **D.** Observed/expected ratio (O/E) of insertion sites in relation to the nine chromatin states defined in *A. thaliana.* Error bars represent the 95% confidence interval obtained by 1000 boots-traps. **E.** GO term analysis of genes containing new *ATCOPIA93* insertions in *HTA9 HTA11* or *hta9 hta11*. **F.** Fraction of essential genes among those targeted by *ATCOPIA93* in *HTA9 HTA11* or *hta9 hta11* F3 seedlings. Statistically significant differences were calculated using the chi-square test. **G.** Density of *ATCOPIA93* insertions over the surface of mouse H2A.Z containing nucleosomes (PDB 1F66). Only 73bp of DNA and associated proteins are viewed down the superhelical dyad and each number (0-7) represents one DNA double helix turn, starting from the central base pair. The exterior circle shows the density of new *ATCOPIA93* insertions per base pair.

### H2A.Z-directed integration is evolutionarily conserved

Unlike *Ty3/Gypsy* LTR-retrotransposons, those of the *Ty1/Copia* superfamily tend to insert within euchromatin (Sultana et al. 2017). To determine if the guiding function of histone H2A.Z is evolutionary conserved, we first focused on *ATCOPIA78*, which is distantly related to *ATCOPIA93* (Figure 5A). There were no private *ATCOPIA78* insertions in the epiRILs, but previous work showed that *ATCOPIA78* can be transcriptionally reactivated by heat stress and mobilized if stressed plants are defective in RNA-directed DNA methylation, such as in the *nrpd1* mutant background (Ito et al., 2011). We therefore assessed the mobilization of *ATCOPIA78* in pools of *nrpd1* seedlings that were subjected or not to heat stress and subsequently grown under normal conditions to produce seeds. One thousand F1 seedlings from each pool were grown under normal conditions and used to perform TE-sequence capture (Figure 5B). A total of 279 *ATCOPIA78* insertions were recovered in the progeny of heat-stressed plants, compared to only two in the progeny of non-stressed plants (Figure 5C). Insertion preferences were similar to those observed with *ATCOPIA93* (Figure 5D, E, F).

**Figure 5.**
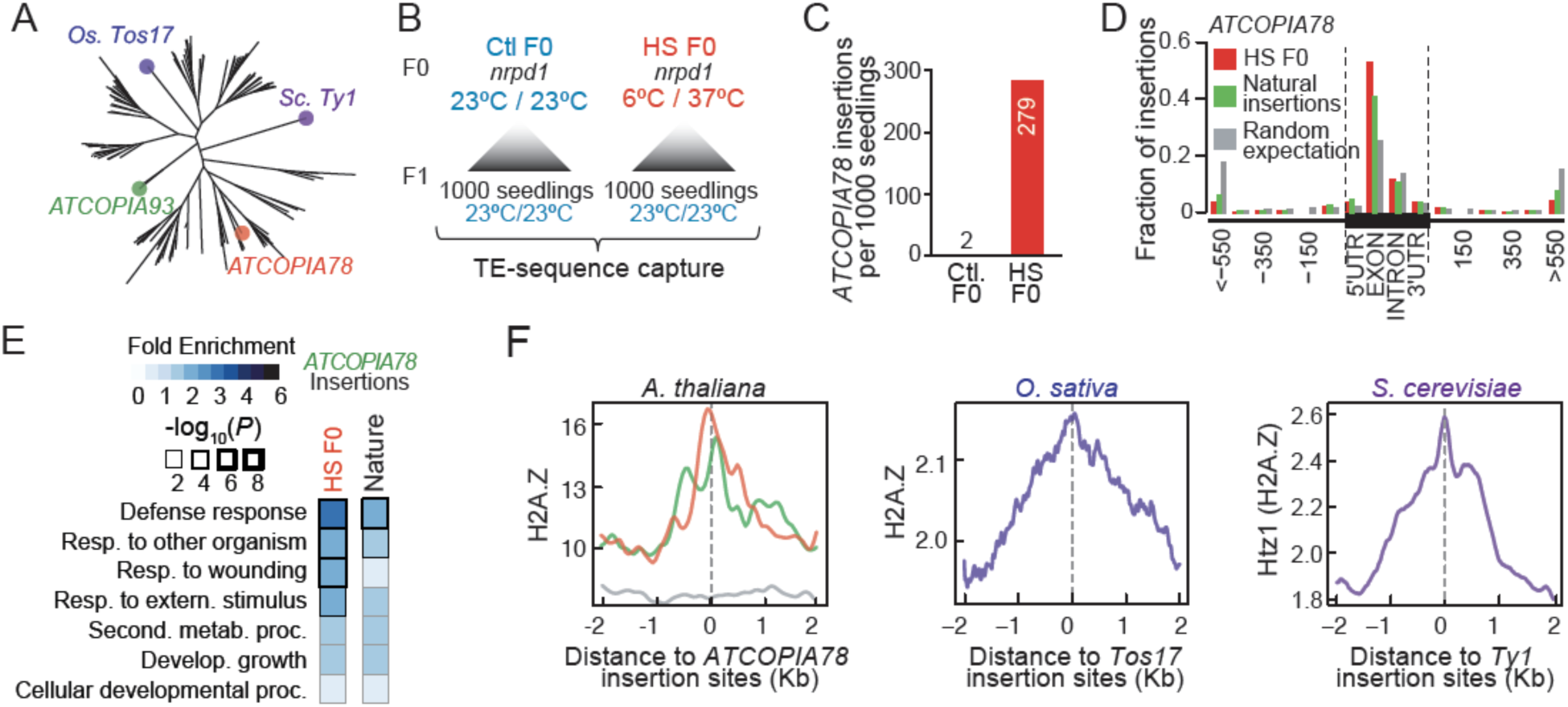
H2A.Z-directed integration of *Ty1/Copia* retrotransposons is evolutionarily conserved. **A.** Phylogenetic analysis of *Ty1/Copia* LTR-retrotransposons from *A. thaliana* as well as *Tos17* and *Ty1* from rice and budding yeast, respectively.
**B.** Experimental strategy for studying transposition landscape in *A. thaliana* of the heat-responsive *ATCOPIA78* LTR-retrotransposon. 1000 *nrpd1* F1 seedlings derived from plants grown under control conditions (Ctl F0) or subjected to heat-stress (HS F0) were subjected to TE-sequence capture. **C.** Number of new insertions detected in Ctl F0 or HS F0. **D.** Metagene analysis showing the distribution of new *ATCOPIA78* insertions detected in heat-stressed *nrpd1* mutant plants (HS F0) or in natural population of *A. thaliana* (Natural insertions). UTR, untranslated transcribed region. **E.** GO term analysis of genes containing new *ATCOPIA78* insertions in heat-stressed *nrpd1* mutant plants (HS F0) or in natural population of *A. thaliana* (Natural insertions). **F.** Metanalysis of *A. thaliana*, rice (*O. sativa*) and yeast (S. *cerevisiae)* H2A.Z levels around *ATCOPIA78, Tos17* and *Ty1* insertion sites, respectively. For *A. thaliana*, experimental and natural insertions are depicted in green and red, respectively.

Unlike for *ATCOPIA93*, numerous recent *ATCOPIA78* insertions were found in nature (Quadrana et al. 2016). Although patterns of integration were similar in the experimental and natural settings (Figure 5D, E and F), the purifying effect of selection was already evident in nature. Specifically, the fraction of natural private insertions within exons was much reduced compared to that found experimentally (Figure 5D). Similarly, none of the 147 recent natural insertions examined were within essential genes, compared to 2.8% of the *ATCOPIA78* insertions detected experimentally (Figure S5). These findings establish a key role for H2A.Z in directing the integration of *COPIA* retrotransposons in nature and further highlight the strong deleterious effects they can cause.

Last, we investigated the integration patterns of *Ty1/Copia* retrotransposons in relation to H2A.Z in a distant plant (rice; Miyao et al., 2003) and in widely divergent species (S. *cerevisiae;* Mularoni et al., 2012). In both cases, experimentally induced insertions were located at sites enriched for H2A.Z (Figure 5F), thus indicating that the guiding role of H2A.Z has been evolutionarily conserved since the last common ancestor of plants and fungi.

### Intronic *ATCOPIA* insertions create epigenetically-inducible alternative transcripts

To assess the functional impact of the strong, chromatin-based insertion biases towards genes observed in the epiRILs, we measured gene expression in six lines by RNA-seq. None of the homozygous *VANDAL21* and *ATENSPM3* insertions had detectable effects on the expression of target genes (Figure 6A), consistent with findings in natural accessions (Quadrana et al. 2016). In marked contrast, six of the 16 distinct homozygous *ATCOPIA93* insertions located within genes affected negatively their expression (Figure 6A). We also performed RNA-seq on one of the six epiRILs taken at a later generation (F16, epiRIL394). At this generation, there were approximately 40 *ATCOPIA93* insertions (counting homozygous insertions as two) per epiRIL and all were silenced and presumably methylated (Figure 6B).

**Figure 6.**
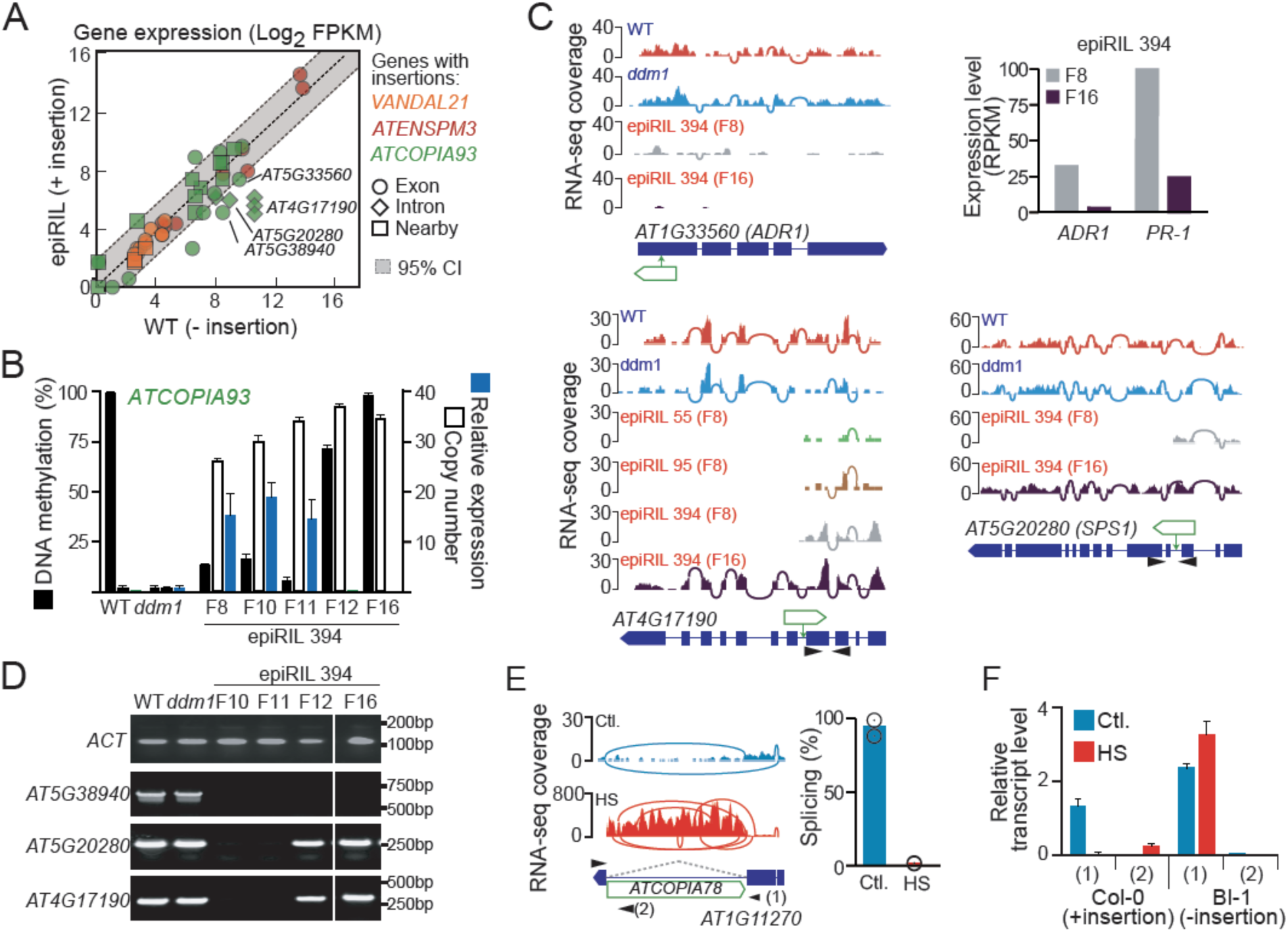
Intronic *Ty1/COPIA* insertions create epigenetically-inducible alternative transcripts. **A.** Expression ratios between epiRILs and wild type plants for genes harboring homozygous *VANDAL21, ATENSPM3* and *ATCOPIA93* insertions. Genes with exonic, intronic or nearby insertions are indicated by circles, boxes or diamonds, respectively. Expected expression ratios and 95% confidence intervals were obtained by sampling 1000 random set of 55 genes and calculating their expression ratio. **B.** q-PCR analyses of DNA methylation, copy number and expression of *ATCOPIA93* in wild type, *ddm1* and epiRIL394, taken at F8 and more advanced generations. Data are mean ± s.d. (n = 2 independent biological experiments). **C.** Genome browser view of RNA-seq coverage over selected genes containing new *ATCOPIA93* insertions. Exon-exon junctions detected by split-reads are represented by arcs that connect exons. Samples containing or lacking TE insertions are highlighted in red and blue, respectively. Expression levels for the gene *ADR1 and PR-1* are shown on the upper right corner. **D.** RT–PCR analysis of genes containing new *ATCOPIA93* insertions in epiRIL394. Primers are indicated by arrow-heads in **B. E.** Genome browser view of RNA seq data for a gene containing an *ATCOPIA78* insertion in Col-0 in plants grown under control conditions (Ctl) or subjected to heat stress (HS). Percentage of splicing for the TE-containing intron (indicated by dashed lines) is shown on the right panel. Data are mean (n = 2 independent biological experiments). **F.** qRT-PCR analyses of *AT1G11270* expression in response to heat stress (HS) in accessions containing (Col-0) or lacking (Bl-1) the intronic *ATCOPIA78* insertion. Primers are indicated in **E** by arrow-heads. Data are mean ± s.d. (n = 2 independent biological experiments).

One *ATCOPIA93* insertion present at F8 and F16 was located in the 5’UTR of gene *AT1G33560,* which encodes an NBS-LRR disease resistance protein, and had a much stronger dampening effect on gene expression at F16 than at F8 (Figure 6C). This result resembles that obtained for another gene in another epiRIL (Marí-Ordóñez et al. 2013). Moreover, reduction in *AT1G33560* expression correlated with down-regulation of *PR-1* (Figure 6C), a gene that is implicated in the response of plants to pathogens and whose expression depends on that of *AT1G33560* (Collier et al., 2011). These findings indicate that epigenetic silencing of newly inserted *ATCOPIA93* insertions can reinforce gene repression with potential phenotypic consequences.

Two *ATCOPIA93*-containing genes, *AT4G17190* and *AT5G20280*, which are implicated respectively in defense against aphids and in nectar secretion (Lin et al. 2014; Bhatia et al. 2015), also showed reduced expression, but only at generation F8. In both cases, *ATCOPIA93* was located within an intron and either in the same or the opposite orientation relative to gene transcription and was associated with transcript truncation at F8, but not at F16 (Figure 6C), when the insertions are epigenetically silent. Examination of DNA methylation, copy number and expression of *ATCOPIA93* as well as expression of the two genes across several generations between F8 and F16 revealed that DNA methylation was established at F12 and coincided with the silencing of all *ATCOPIA93* copies (Figure 6B) as well as the proper splicing of the insertion-containing intron (Figure 6D). These results demonstrate that the deleterious effects of intronic *ATCOPIA93* insertions can be alleviated once they become epigenetically silent.

We last investigated a natural intronic *ATCOPIA78* insertion that is present in the reference Col-0 genome but absent from other accessions (Figure S6). Analysis of publicly available RNA-seq data (Pietzenuk et al. 2016) indicated that the insertion is epigenetically reactivated by heat stress and that reactivation is associated with the production of a truncated transcript from the target gene (Figure 6E). To demonstrate causality, we compared the splicing level of the second intron of the target gene (*AT1G11270*) in Col-0 plants subjected or not to heat stress as well as in plants of the Bl-1 accession, which does not contain any *ATCOPIA78* insertion within this gene. While the second intron was spliced in heat-stressed and control Bl-1 plants, this was not the case for Col-0 plants, which showed splicing only under non-stress conditions (Figure 6F). Given that *ATCOPIA78* is among the most active TE families in nature and that its activity correlates with several geo-climatic variables (Quadrana et al. 2016), it is likely that numerous natural *ATCOPIA78*-containing alleles of genes are similarly endowed with the possibility to modulate their expression in response to environmental stress. Whether such alleles confer selective advantages remains to be determined.

## Discussion

### TEA lines: when mobile TEs are caught in the act

Whole genome sequencing of MA lines has been extremely valuable in determining the rate, spectrum and genome-wide distribution of spontaneous point and other small-size mutations in several model organisms (Ossowski et al. 2010; Zhu et al. 2014; Denver et al. 2009; Keightley et al. 2009). Results of these experiments indicated that small-size mutations occur almost randomly throughout the genome, although local modulations were observed, notably in relation to certain DNA sequences, recombination, transcription activity, and chromatin marks, including DNA methylation (Makova and Hardison 2015). However, MA lines did not provide much information about the heritable mutational landscape generated by TE mobilization, mainly because transposition is typically rare and also because non-reference TE insertions are difficult to detect using standard short-read sequencing strategies.

Based on a unique experimental system of *A. thaliana* epiRILs, in which transposition activity was kick-started but that otherwise resembled MA lines, we documented >8000 new TE insertions. These were produced mainly by three TEs, which belong to two DNA transposon (*VANDAL21* and *ATENSPM3*) and one LTR-retrotransposon (*ATCOPIA93*) families that are among the most active in nature (Quadrana et al. 2016). Furthermore, our modeling indicated that the insertion mutations produced by each of these three TEs in the epiRILs likely accumulated following a transposition-genetic drift scenario. Thus, the epiRILs can be defined operationally as TEA lines and as such they provide a first comprehensive assessment of the actual rate and genome-wide distribution of heritable mutations generated by TE mobilization in any species.

Using the epiRILs and additional experimental populations, we showed that *VANDAL21*, *ATENSPM3, ATCOPIA93* as well as another member (*ATCOPIA78*) of the *Ty1/Copia* superfamily of LTR-retrotransposons preferentially integrate within or close to three distinct sets of genes, each characterized by specific chromatin features. Moreover, we obtained strong evidence that *ATENSPM3* and the two *ATCOPIA* families generate highly deleterious mutations that are immediately purged by natural selection.

Collectively, our findings provide a first comprehensive experimental demonstration that mutations generated by TEs have radically distinct properties than the spontaneous small size mutations documented in MA lines. First, because of strong TE-specific integration preferences linked to chromatin, TE-caused mutations are distributed non-uniformly across the genome, with the implication also that their repertoire may vary substantially for any given TE as a result of changes in chromatin states, such as the loss of heterochromatin. Second, because many TEs target chromatin states that are associated with genes, their insertion, as well as their excision in the case of DNA transposons, tend to have drastic effects on gene expression. This is however not an obligate outcome, as exemplified by the lack of any discernible consequences of *VANDAL21* integration on gene transcription. Third, the functional impact of TE insertions can vary in a reversible manner over very short times, through their epigenetic silencing, which can be influenced by the environment, as in the case for *ATCOPIA78*. Last, TE insertions accumulate discontinuously, as a result of episodic, TE-specific reactivation, and at rates that may differ widely between TEs and environments.

### Retrotransposition-driven allelic heterogeneity associated with adaptive traits

Although WGS has revealed that TEs are powerful engine of genome as well as organism evolution (Chuong, Elde, and Feschotte 2016), we still lack a clear understanding of the impact of their mobilization within any given species. Our finding that LTR-retrotransposons belonging to the *Ty1/Copia* superfamily preferentially integrate within environmentally responsive genes provide valuable information in this respect. As we have also shown previously that mobilization of diverse *ATCOPIA* elements is associated with climate (Quadrana et al. 2016), it is tempting to speculate that this superfamily of TEs facilitates adaptation to changing or local environments. Consistent with this hypothesis, disease resistance genes are characterized by a high load of *COPIA* insertions and extensive allelic heterogeneity in nature (Quadrana et al, 2016; Kawakatsu et al., 2016). Thus, recurrent retrotransposition within these genes may play a critical role in the rapid evolution and expansion of the innate immune system in plants. We have demonstrated however that most *ATCOPIA* insertions have severe deleterious effects, athough some of these effects may be mitigated or fully erased once the insertions are epigenetically silenced. This mitigation can be environmentally dependent, thus endowing TE-containing alleles with unique properties in fluctuating environments (Figure 7A).

**Figure 7.**
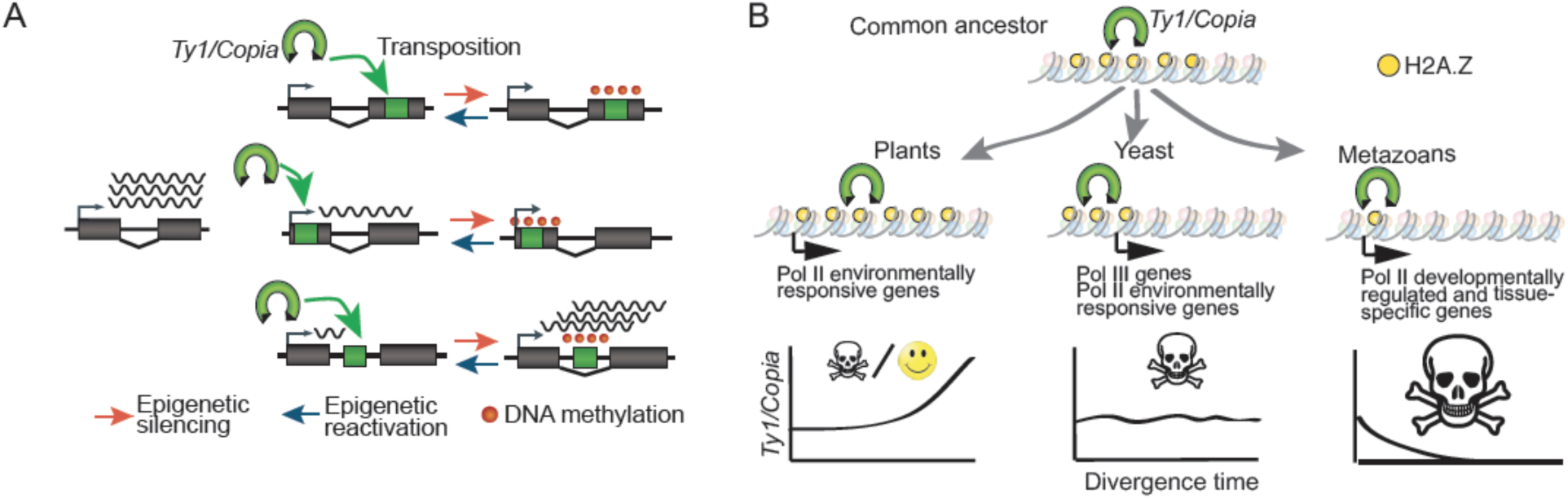
A model to depict the functional consequences of new TE insertions and the evolutionary fate of *Ty1/Copia* retrotransposons in different organisms. **A.** *De novo* retrotransposon insertions within genes impact their expression in multiple ways. While retrotransposition within internal exons systematically disrupt gene expression, insertions in 5’ regions or introns have an immediate dampening effect, which can be either aggravated or mitigated following epigenetic silencing of the inserted TE, respectively. Note that DNA methylation is depicted as spreading from the *COPIA* insertion into adjacent sequences, which remains to be determined experimentally. **B.** H2A.Z-guided integration of *Ty1/Copia* is ancestral. Functional diversification of H2A.Z between kingdoms determined the preferential integration of *Ty1/Copia* towards fast evolving genes in plants and yeast and towards developmentally regulated genes in animals. This differentiation may explain the high and low invasions success of *Ty1/Copia* retrotransposons in plants and animals, respectively.

### H2A.Z and the different fate of *Ty1/Copia* retrotransposons in plants, yeast and animals

TEs need to keep moving in order to prevent their demise by the accumulation of inactivating mutations. However, because uncontrolled mobilization compromises host survival and that of TEs, different mechanisms have evolved that limit TE activity, notably epigenetic silencing, or that target integration to reduce the mutational load they generate (Sultana et al. 2017). For example, the LTR-retrotransposon *Ty5* from yeast interacts with the heterochromatic factor silent information regulator 4 (Sir4), which directs *Ty5* integration within gene-poor regions (Sultana et al. 2017). Here, we have uncovered another targeting mechanism based on the histone variant H2A.Z, which is among the most conserved histone H2As (Henikoff and Smith 2015). H2A.Z guides the integration of *Ty1/Copia* elements not only in *A. thaliana* but presumably also in rice and yeast. Furthermore, as in plants, the *S. cerevisiae Ty1* retrotransposon integrates preferentially within the arrays of H2A.Z-containing nucleosomes that are located upstream of RNA polymerase III (Pol III)-transcribed genes (Figure 5F; Albert et al., 2007; Baller et al., 2012), aided by the Pol III subunit AC40 (Bridier-Nahmias et al. 2015). Disruption of the AC40-integrase interaction leads to a redistribution of *Ty1* insertions towards subtelomeres (Bridier-Nahmias et al. 2015), which are enriched in environmentally responsive genes (Brown, Murray, and Verstrepen 2010). As in plants, these genes are enriched in H2A.Z-containing nucleosomes (Meneghini, Wu, and Madhani 2003; Sadeghi et al. 2011; Albert et al. 2007). These findings suggest therefore that AC40 serves to further restrict the integration of *Ty1* to a subclass of genes containing arrays of H2A.Z, which otherwise would integrate preferentially within or around responsive genes.

In animals, H2A.Z is essential and mainly found at developmentally regulated genes, where it occupies nucleosomes that flank the promoter (Henikoff and Smith 2015). Given the conserved role of H2A.Z in guiding the integration of *Ty1/Copia* retrotransposons, their mobilization should therefore have catastrophic effects in animals, which may explain why this superfamily of retrotransposons is virtually absent from animal genomes (Huang, Burns, and Boeke 2012). In other words, functional diversification of H2A.Z between plants and animals may have directed the opposite fate of *Ty1/Copia* retrotransposons in these two kingdoms (Figure 7B). This in turn opens up the possibility that the evolutionary fate of other TEs could also be explained by similar chromatin-directed integration biases.

## Acknowledgments

We thank members of the Colot group and especially Pierre Baduel for discussions and critical reading of the manuscript and S. Khouider for help with plant crosses. We thank the Genomics and Informatics facilities at IBENS for their help. Support was from the Agence National de la Recherche (ANR-09-BLAN-0237 to V.C. and P.W.), the Investissements d’Avenir ANR-10-LABX-54 MEMO LIFE, ANR-11-IDEX-0001-02 PSL* Research University (to V.C.), the European Union Seventh Framework Programme Network of Excellence EpiGeneSys (Award 257082, to V.C.) and the Centre National de la Recherche Scientifique (MOMENTUM program, to L.Q.). M.E. was supported by a PhD studentship from the French Ministry of Research and by a postdoctoral fellowship from MEMOLIFE. Initial support for L.Q. was provided by MEMOLIFE and EpiGeneSys. V.B. was supported by a PhD studentship from PSL Research University and from MEMOLIFE.

## Author contributions

VC, PW and JMA conceived the project, with substantial additional input from LQ. ME and EC performed DNA and RNA extraction; JG, SE and JMA produced the WGS data; AG, ME, JMA and LQ performed the detection of TE insertions in the epiRILs; VB identified excision footprints in the epiRILs. LQ performed all of the other bioinformatic analyses, as well as the TE sequence capture and RT-QPCR experiments. LQ and AS performed the heat-stress experiments. LQ and VC interpreted the results. LQ and VC wrote the manuscript.

## MATERIALS AND METHODS

### Experimental model and subject details

The following *A. thaliana* plants were used: wild type Col-0 and Bl-1 accessions. The Col-0 *ddm1-2* mutant and the epiRILs population (Johannes et al. 2009), the Col-0 *nrpd1-3* mutant (H. Ito et al. 2011) and the Col-0 *hta9-1 hta11-2* double mutant (March-Díaz et al. 2008). Unless stated otherwise, all plants were grown in long-days (16h:8h light:dark) at 23°C.

### Genomic DNA sequencing using mate-pairs

Genomic sequencing was performed as described before (Gilly et al, 2014). Briefly, DNA was extracted from 10-20 seedlings grown under long day conditions, using DNeasy Qiagen kits. About 10 μg of genomic DNA was sonicated to a 4-6 kb size range using the E210 Covaris instrument (Covaris, Inc., USA). Libraries were prepared following Illumina’s protocol (Illumina Mate Pair library kit), starting with size-selected (approximately 5kb) fragments, which were end-repaired, biotin labeled and circularized. Linear DNA was eliminated by digestion and circularized DNA was fragmented to 300-700 bp using the E210 Covaris. Biotinylated DNA junctions were purified using streptavidin, end-repaired and 3’-adenylated in order to ligate Illumina adapters. Junction fragments were PCR-amplified using Illumina adapter-specific primers and amplified fragments within the 350-650 bp size range were selected for sequencing. Each library was sequenced using 100 base-length read chemistry in a paired-end flow cell on the Illumina GAIIx (2 lanes) or HiSeq2000 (1 lane) (Illumina, USA).

### Mapping and detection of TE insertions using WGS

Reads were mapped with BWA v.0.6.1 using the parameters -R 10000 -l 35 -O 11, and the parameters n 10000 N 10000 -s for sampe, onto the TAIR10 reference sequence. Reads hanging over chromosome ends were removed using picard CleanSam and duplicate pairs were removed using picard MarkDuplicates. TE insertions were detected by implementing TE-Tracker software (available at http://www.genoscope.cns.fr/TE-Tracker) exactly as described before (Gilly *et al.*, 2014). WGS did not produce sufficient coverage (<10X) for 16 of the 123 epiRILs analyzed at generation F8, and these 16 epiRILs were not considered further (Table S1). TE-Tracker is a computational method that we have previously developed for accurately detecting both the identity and destination of newly mobilized TEs in genomes re-sequenced using mate-pair libraries (Gilly et al, 2014). Importantly, TE-Tracker does not rely on prior annotation, yet is able to integrate it, making the results easily interpretable. Briefly, TE-Tracker uses paired reads mapping information to identify discordant reads that mapped partially over TE-sequences to detect the position of TE insertions. Insertion site positions were refined at the single nucleotide resolution by exploiting sequence information contained in split-reads. To this end, we implemented the software SPLITREADER (available at https://aithub.com/LeanQ/SPLITREADER: Quadrana et al., 2016). Homozygous and heterozygous insertions were defined based on the normalized number of reads supporting each insertion event (Table S2). In addition, this approach enabled us also to identify insertions that were likely present in only one of the 10-20 seedlings used to extract DNA and which reflect transposition during the reproductive phase of the parent. These insertions were also called heterozygous, as this was likely the case and to reflect their very recent ancestry. Conversely, our approach was designed to exclude poorly supported insertions, which could reflect either mapping artifacts or rare somatic events. Finally, visual inspection was carried out for a random sample of over 200 insertion events and their homozygous or heterozygous status was confirmed in each case.

### TE-sequence capture

TE sequence capture was performed on exactly 1000 seedlings in all cases except for the F3 progeny of *hta9 hta11* line 1, where only 477 seedlings were recovered (see main text and Figure 4A and 5B for details of the plant materials used). Seedlings were grown under control (long-day) conditions and genomic DNA was extracted using the CTAB method (Murray and Thompson 1980). In order to assess the sensitivity of TE-sequence capture, we added 1ng of genomic DNA extracted from epiRILs 394 (generation F16) to 1ug of genomic DNA extracted from the 477 F3 seedling of *hta9 hta11* line 1 prior to library construction (i.e. 1:1000 dilution of the spiked-in genomic DNA). Libraries were prepared using 1*μ*g of DNA and TruSeq paired-end kit (Illumina) following manufacturer instructions. Libraries were then amplified through 7 cycles of ligation-mediated PCR using the KAPA HiFi Hot Start Ready Mix and primers AATGATACGGCGACCACCGAGA and CAAGCAGAAGACGGCATACGAG at a final concentration of 2*μ*M. 1*μ*g of multiplexed libraries were then subjected to TE-sequence capture exactly as previously reported (Quadrana et al. 2016). Enrichment for captured TE sequences was confirmed by qPCR and estimated to be higher than 1000 fold. Pair-end sequencing was performed using one lane of Illumina NextSeq500 and 75bp reads. About 42 million pairs were sequenced per library and mapped to the TAIR10 reference genome using Bowtie2 v2.3.2 (Langmead and Salzberg 2012) with the arguments --mp 13 --rdg 8,5 --rfg 8,5 --very-sensitive. An improved version of SPLITREADER (available at https://github.com/LeanQ/SPLITREADER) was used to detect new TE insertions. Briefly, split-reads as well as discordant reads mapping partially on the reference sequence of *ATCOPIA93* and *ATCOPIA78* (obtained from RepBase update) were identified, soft clipped and remapped to the TAIR10 reference genome using Bowtie2 (Langmead and Salzberg 2012). Putative insertions supported by at least one split- and/or discordant-reads at each side of the insertion sites were retained. Insertions spanning centromeric repeats or coordinates spanning the corresponding donor TE sequence were excluded. In addition, putative TE insertions detected in more than one library were excluded to retain only sample-specific TE insertions (Table S3). More than 80% of new TE insertions present in epRIL394 F16 were detected, confirming that the sensitivity of our TE-sequence capture and computational approach is higher than 1:1000. In addition, no non-reference insertions were detected in 1000 F1 seedlings of wild type Col-0, highlighting the specificity of our approach.

### Detection of Tos17 non reference insertions in rice genomes

A total of 16,784 *Tos17* non-reference flanking sequences (Miyao et al. 2003) were retrieved from GeneBank and mapped to the reference rice genome using Minimap2 v2.11-r797 (H. Li 2018), which enabled us to identify 14,258 insertion points with high confidence (Table S4).

### Detection of mutations within transposed copies

Discordant mate-pair reads mapping within a 6kb interval either upstream or downstream of each insertion site were extracted and re-mapped using Bowtie2 (Langmead and Salzberg 2012) over a library constructed with the specific donor TE sequence only. Sequence variants were detected using samtools mpileup V1.2.1 and only variants with a quality of at least 30 were kept. Long deletions were detected as regions without coverage and breakpoints were reconstructed by local assembly using Velvet V1.2.09 (Zerbino and Birney 2008).

### Analysis of global and local enrichment of new TE insertions

To assess if new TE insertions are enriched in pericentromeric regions, their number within these regions was compared with that expected from a random distribution. The expected distribution was calculated by randomizing 10^4^ times the position of new TE insertions across the chromosomes (genomic regions showing coverage deviation, the inner pericentromeres, or coordinates spanning the corresponding donor TE sequence were excluded). This set of random positions was used as a control for all subsequent analyses. Insertion distribution over wild-type- and *ddm1*-derived regions was obtained by counting the number of new TE insertions within intervals delimited by at least two consecutive stable hypermethylated or hypomethylated regions, respectively (Cortijo et al. 2014). Overrepresentation over genes and neighboring sequences was performed using a meta-gene analysis. Briefly, protein coding gene features were extracted from the TAIR10 annotation and coordinates of non-reference TE insertions with TSDs were crossed with the set of genic features according to the following stepwise hierarchy: 5’ UTR > 3’ UTR> exon > intron > intergenic regions. For insertions that do not overlap protein-coding genes, the distance to the closest gene was calculated and reported as negative or positive distance according to the gene orientation. Overrepresentation over chromatin states was performed by comparing the number of new TE insertions and randomly generated TE insertions located within each chromatin domain (Sequeira-Mendes et al. 2014). Density of ATCOPIA93 insertions were obtained by calculating the distance between insertion sites and the middle point of the nearest well positioned nucleosome mapped in Col-0 (Table S5; Lyons and Zilberman, 2017). Gene ontology (GO) analyses were performed using AGRIGO (http://bioinfo.cau.edu.cn/agriGO/) and as input the ID of genes that contain a TE insertion within the limits of their annotation.

### Analysis of chromatin features at insertion sites

4kb regions centered around insertion sites were defined and used to extract normalized coverage of DnaseI hypersensitivity (Sullivan et al. 2014), Mnase accessibility (G. Li et al. 2014), H3K27me3 (C. Li et al. 2015), H2A.Z enrichment level (Coleman-Derr and Zilberman 2012) and well-positioned nucleosomes (Lyons and Zilberman 2017). The same approach was used for the analysis of H2A.Z enrichment in rice (Zahraeifard et al. 2018) and of htz1 from yeast (Gu et al. 2015). Average normalized coverage was then calculated for each bp and plotted using the *smooth.spline* function in R.

### epiQTL mapping of transposition activity

Using TE copy number as phenotype and a total of 126 parental differentially methylated regions (DMRs) that segregate in a Mendelian fashion in the epiRILs (i.e. stable DMRs) as physical markers (Cortijo et al. 2014), we implemented the multiple QTL model (mqmsacn) from the R/qtl package. Genome-wide significance was determined empirically for each trait using 1000 permutations of the data. LOD significance thresholds were chosen to correspond to a genome-wide false positive rate of 5%.

### Transposition-drift modeling of insertion accumulation

In the absence of selection, TE invasion is mostly determined by the rate of transposition and rate of fixation of insertions by random segregation (i.e. drift). Thus, we constructed individual-based transposition-drift models, all starting with an initial number of copies all equally active (with a rate *K* of new copies per generation) and with each new copy being also equally active. New copies arise in the heterozygous state and can be inherited following a Poisson distribution according to Mendelian segregation. The model also considers TE elimination by excision, which occurs at rate *E* per transposition event. Additionally, concerted silencing of all active copies may occur when copy number reach the threshold *l*. Simulations were run 1000 times using a wide space of parameter values (*K*={0,0.1,…,1}, *E*={0,0.2,…,1} and *I*={20,21,…,60}. Distribution of homozygous and heterozygous insertions between simulated and observed data at F8 was evaluated using a two-dimensional goodness-of-fit test.

### Transcriptome analysis

RNA from wild type, *ddm1*, epiRIL55 (F8), epiRIL95 (F8), epiRIL260 (F8), epiRIL439 (F8), epiRIL454 (F8) epiRIL394 (F8) and epiRIL394 (F16) was isolated using Rneasy Plant Minikit (Qiagen) according to the supplier’s instructions. Contaminating DNA was removed using RQ1 DNase (Promega). One μg of total RNA was processed using TruSeq Stranded Total RNA kit (Illumina) according to the supplier’s instructions. About 20M 76nt-long single-end reads were obtained per sample on the Illumina HiSeq2000. Expression level was calculated by mapping reads using STAR v2.5.3a (Dobin et al. 2013) on the *A. thaliana* reference genome (TAIR10) with the following arguments --outFilterMultimapNmax 50 --outFilterMatchNmin 30 -- alignSJoverhangMin 3 --alignIntronMax 10000. Duplicated pairs were removed using picard MarkDuplicates. Counts were normalized and annotations were declared differentially expressed between samples using DESeq2 (Love, Huber, and Anders 2014). When specified, uniquely mapped reads (mapping quality > 10) were selected with samtools (H. Li et al. 2009). RNAseq data from heat-stressed plants were obtained from (Pietzenuk et al. 2016) and analyzed as described above. Splicing of intronic TE insertions was calculated as described previously (Teixeira et al. 2017). Briefly, the number of split-reads (SR) and non-split-reads (NSR, which should be fully and uniquely contained within the interval surrounding the same exon–intron junction) mapping to an exon-intron junction was extracted and the ratio SR/(SR+NSR)*100 was then calculated.

### Quantification of expression level, copy number and DNA methylation

RNA was extracted using the RNeasy plant mini kit (Qiagen) from Col-0 or Bl-1 plants grown under normal conditions (10 days old seedlings grown in liquid medium) or subjected to heat shock treatment (H. Ito et al. 2011). RT-qPCR was performed as described previously (Quadrana et al. 2016). Primer sequences are provided in Table S6. RT-qPCR results (two biological replicates) are indicated relative to those obtained for a gene (*AT5G13440*) that shows invariant expression under multiple conditions. Copy number and DNA methylation of *ATCOPIA93* was performed as described before (Marí-Ordóñez et al. 2013).

### Data and software availability

Sequencing data has been deposited in the European Nucleotide Archive (ENA) under project XXXXX.

---

**Figure S1.**
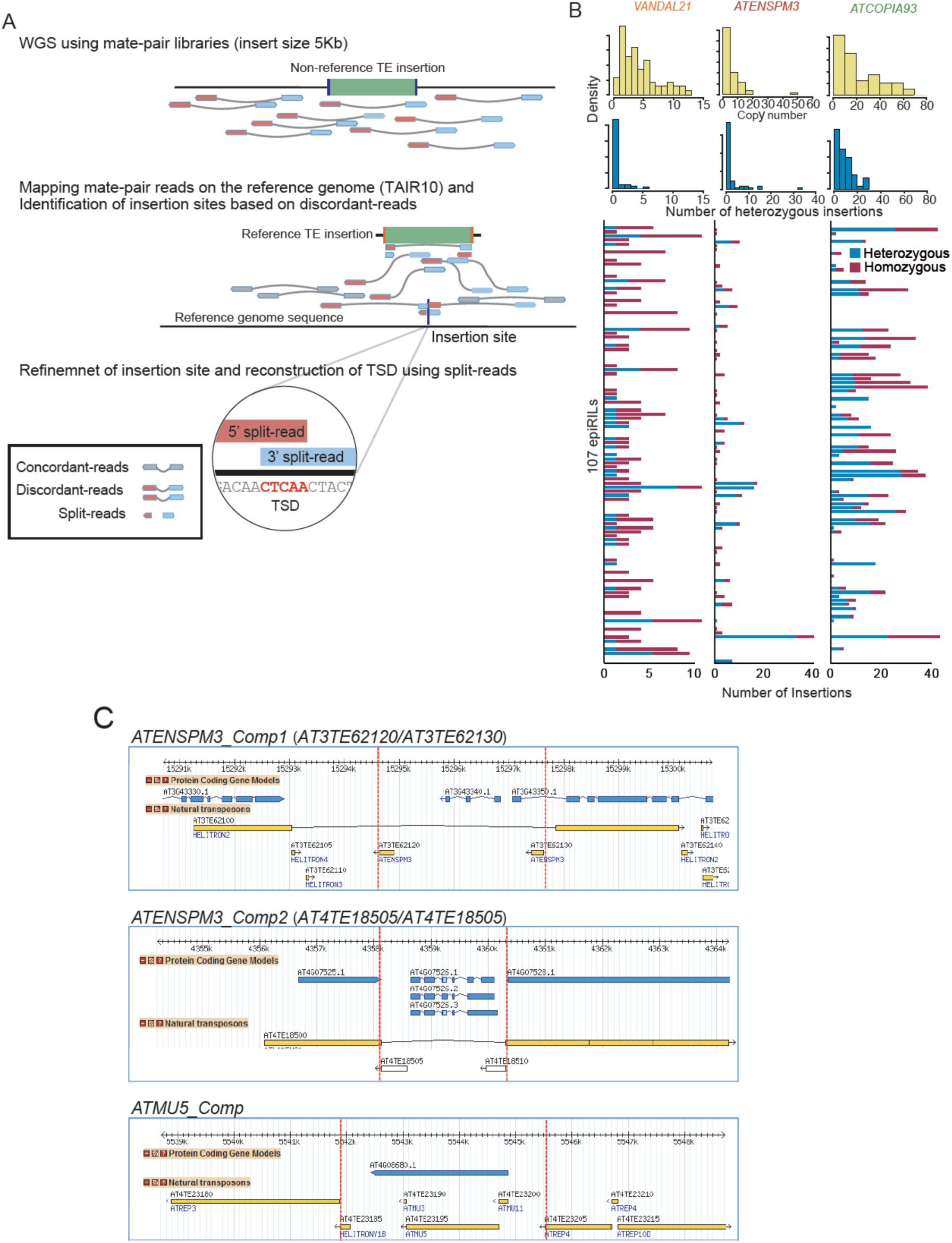
Identification of new TE insertions in epiRILs. **A.** Schematic representation of the bioinformatic method used to detect new TE insertions. Sequenced reads around a newly inserted TE-copy (top half) produce discordant read mappings when aligned with the reference sequence (bottom half). Dashed light-red and light-blue arrows represent the mate-pairs reads linking the left and right extremities of the insertion breakpoint with the donor TE sequence. **B.** The top two rows of panels indicate the distribution of the number of private insertions (top: all; bottom: heterozygous only) among epiRILs for each TE family. The number of homozygous and heterozygous private insertions in each epiRIL (ordered by name) is indicated below for each of the three TE families. **C.** Genome browser view of the three composite mobile TEs (delineated by the dotted red lines) identified in the present study.

**Figure S2.**
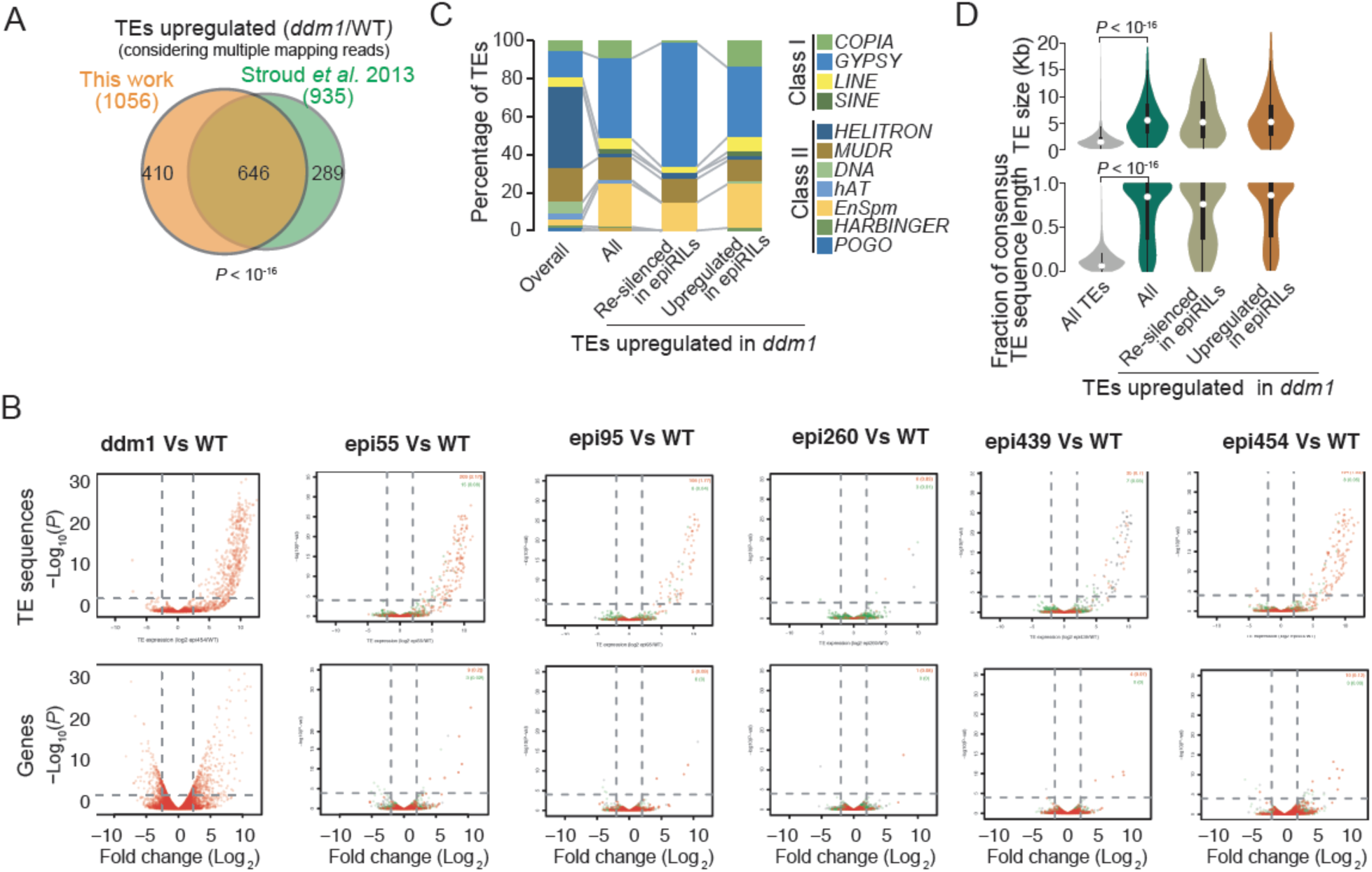
Sustained transcriptional activation of TEs does not associate systematically with transposition. **A.** Comparison of the number of TE sequences upregulated in siblings of the *ddm1* parent used to generate the epiRILs population with a that obtained previously (Stroud et al, 2013). Statistical significance of the overlap was obtained using the Chi-square test. **B.** Differential expression analysis of TEs and genes between wild type and *ddm1*as well as five epiRILs. Annotations located in wild-type- or *ddm1*-detived regions are indicated by green and red dots, respectively. **C.** Identity of the TE sequences upregulated in *ddm1* and either re-silenced or stably upregulated in the five epiRILs. **D.** Distribution of sequence lengths and fraction of TE consensus length of TEs upregulated in *ddm1* and either re-silenced or stably upregulated in the five epiRILs.

**Figure S3.**
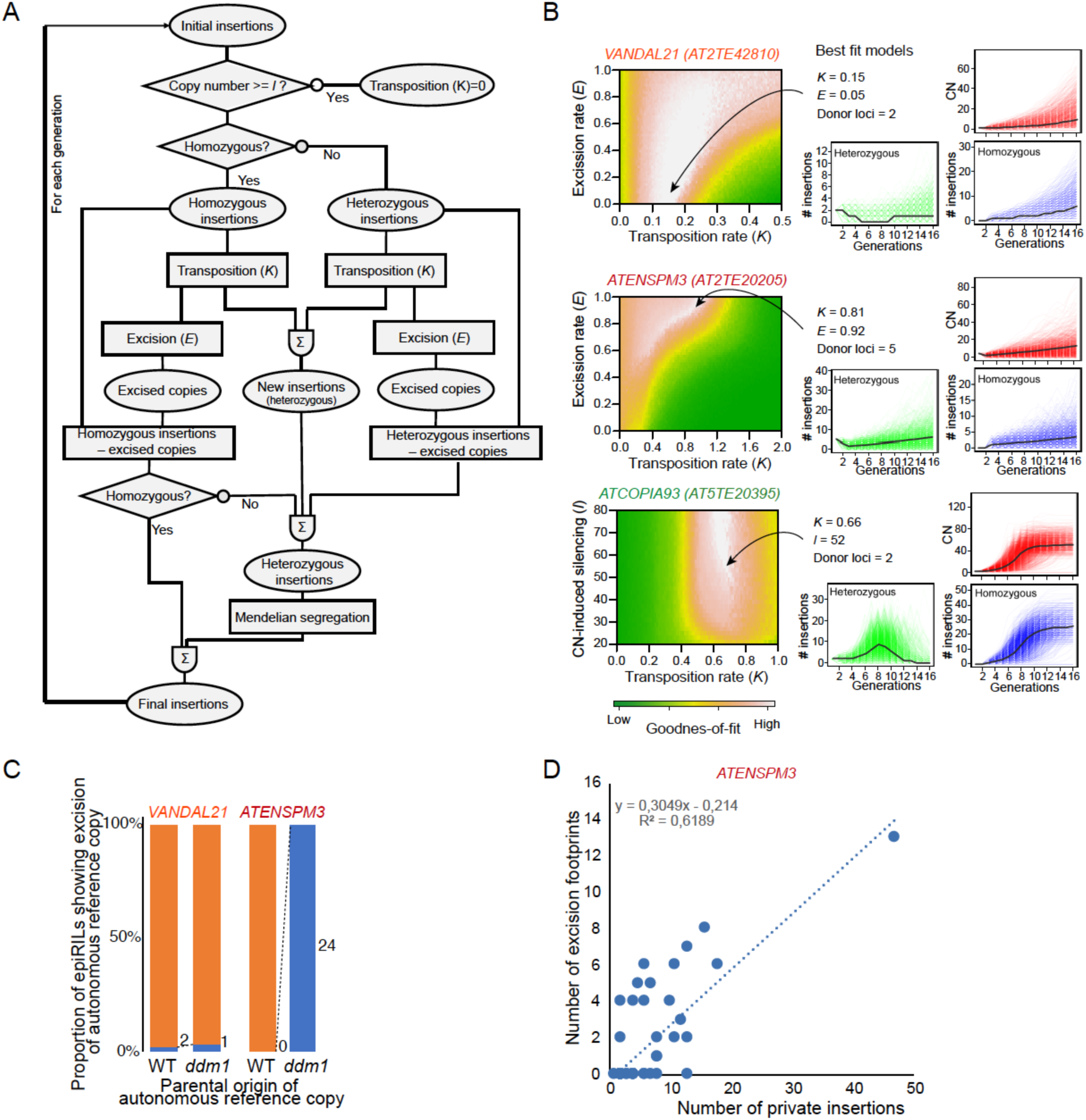
Modeling of insertion accumulation dynamics. **A.** Schematic representation of the transposition-drift model developed to reconstruct transposition dynamics. **B.** Goodnessoffit between simulated and observed data. Patterns of insertion accumulation produced by the best fitted models. **C.** Proportion and number of epiRILs showing excision of the autonomous reference copy for *VANDAL21* and *ATENSPM3* in relation to their parental origin. **D.** Correlation between the number of private *ATENSPM3* insertions and excision footprints detected in the epiRILs.

**Figure S4.**
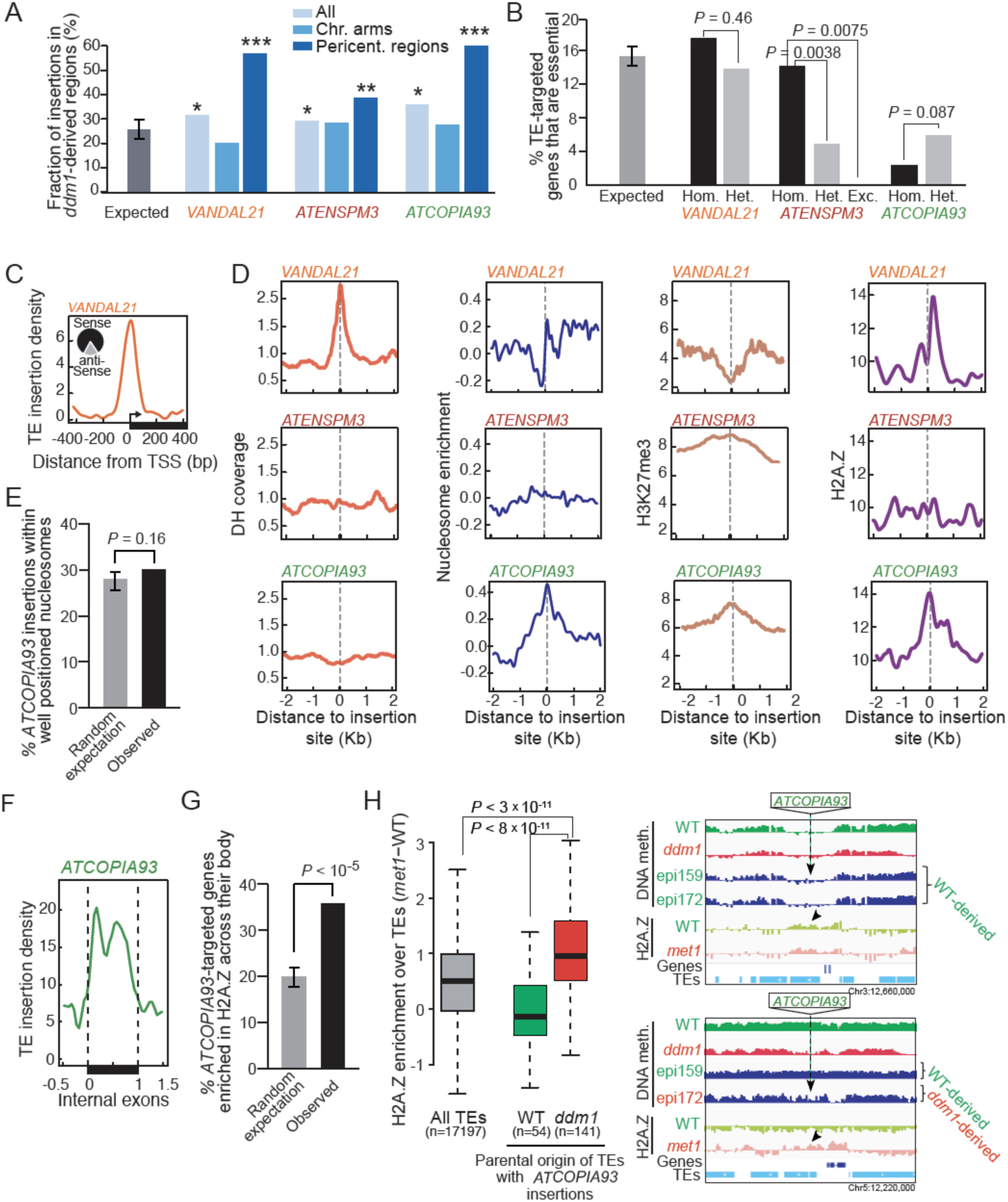
*ddm1*-derived pericentromeric regions are preferentially target by TE insertions. **A.** Proportion of private TE insertions in *ddm1*-derived intervals in relation to their chromosomal position. Statistically significant differences compared to the expected values are indicated (Chi-square test,* P <0.05, ** P < 0.001, *** P<0.0001). **B.** Fraction of essential genes among those targeted by *VANDAL21, ATENSPM3* or *ATCOPIA93* in the epiRILs. Fraction of essential genes among those containing short indels compatible with *ATENSPM3* excision footprints is also indicated. Statistical significance for each comparison was obtained using the Chi-square test. **C.** Density of *VANDAL21* insertions around transcriptional start sites (TSS). The fraction of insertions that are in the same (sense) or opposite (antisense) orientation relative to the targeted gene are indicated. **D.** Meta-analysis of of DNAse hypersensitivity (DH) levels as well as nucleosome, H3K7me3 and H2A.Z densities around insertion sites for *VANDAL21, ATENSPM3* and *ATCOPIA93*. **E.** Proportion of *ATCOPIA93* private insertions within well-positioned nucleosomes. Expected distribution was obtained by randomizing 1000 times insertion site positions across the genome and performing a randomization test. Errors bars represent the 95% confidence interval. **F.** Density of *ATCOPIA93* insertions within internal exons of protein coding genes. **G.** Fraction of genes containing *ATCOPIA93* private insertions that are also enriched in H2A.Z across their body. Expected distribution was obtained by randomizing 1000 times insertion site positions across the genome and performing a randomization test. Errors bars represent the 95% confidence interval. **H.** Relative level of H2A.Z in *met1* compared to wild type (Zilberman et al 2008) over all TEs (grey box) and in the subset of TEs that contain *ATCOPIA93* insertions in the epiRILs (green box: TEs located in wild-type-derived regions; red box: TEs located in *ddm1*-derived regions (in green and red, respectively). Genome browser views of DNA methylation and H2A.Z over TEs containing *ATCOPIA93* insertions within wild-type- and *ddm1*-derived intervals (top and bottom panel, respectively) are depicted on the right. Insertion sites are indicated by an arrow and samples showing hypomethylation across the region are highlighted in red. Statistical significance for each comparison were obtained by Mann–Whitney test.

**Figure S5.**
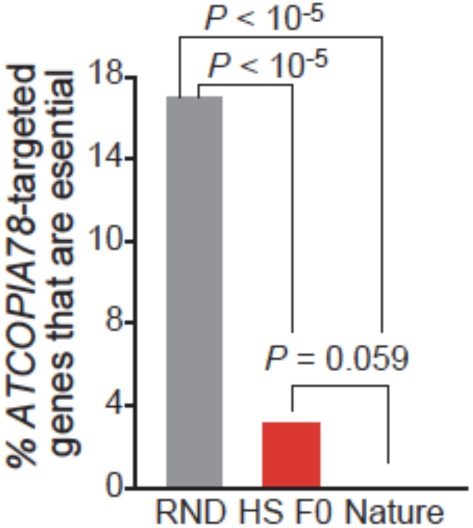
ATCOPIA78 avoids integration within essential genes. **A.** Fraction of essential genes among those targeted by *ATCOPIA78* in significance for each comparison was obtained using the Chi-square test.

**Figure S6.**
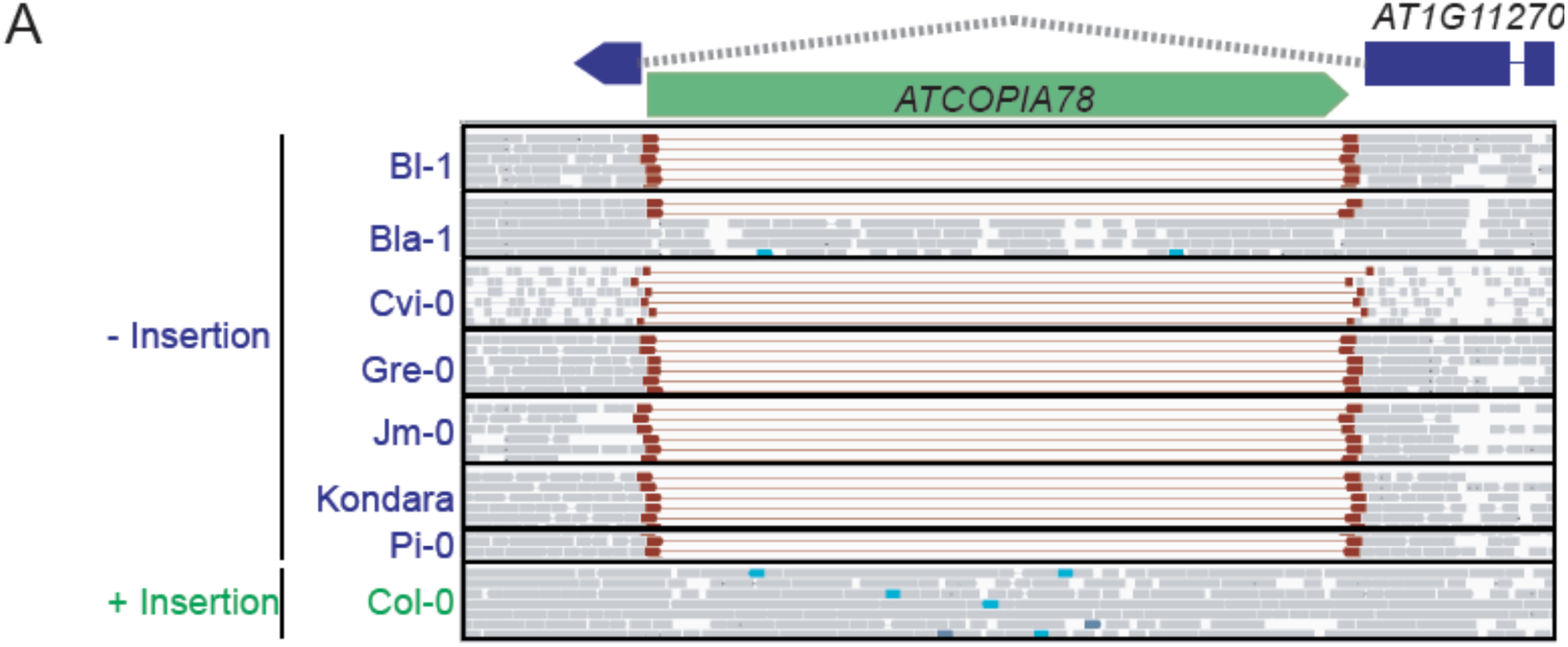
Col-0 allele of *AT1G11270* contains a recent *ATCOPIA78* insertion in the second intron. **A.** Genome browser view of sequence reads produced by WGS over *AT1G11270* for Col-0, which contains an *ATCOPIA78* insertion within the second intron, and seven other accessions that lack this insertion.

## REFERENCES

Albert, Istvan, Travis N. Mavrich, Lynn P. Tomsho, Ji Qi, Sara J. Zanton, Stephan C. Schuster, and B. Franklin Pugh. 2007. “Translational and Rotational Settings of H2A.Z Nucleosomes across the Saccharomyces Cerevisiae Genome.” Nature 446 (7135): 572–76. https://doi.org/10.1038/nature05632.

Baller, Joshua A., Jiquan Gao, Radostina Stamenova, M. Joan Curcio, and Daniel F. Voytas. 2012. “A Nucleosomal Surface Defines an Integration Hotspot for the Saccharomyces Cerevisiae Ty1 Retrotransposon.” Genome Research 22 (4): 704–13. https://doi.org/10.1101/gr.129585.111.

Bhatia, Varnika, Jaya Maisnam, Ajay Jain, Krishan Kumar Sharma, and Ramcharan Bhattacharya. 2015. “Aphid-Repellent Pheromone E-β- Farnesene Is Generated in Transgenic Arabidopsis Thaliana over-Expressing Farnesyl Diphosphate Synthase2.” Annals of Botany 115 (4): 581–91. https://doi.org/10.1093/aob/mcu250.

Bridier-Nahmias, Antoine, Aurélie Tchalikian-Cosson, Joshua A. Baller, Rachid Menouni, Hélène Fayol, Amando Flores, Ali Saïb, Michel Werner, Daniel F. Voytas, and Pascale Lesage. 2015. “An RNA Polymerase III Subunit Determines Sites of Retrotransposon Integration.” Science (New York, N.Y.) 348 (6234): 585–88. https://doi.org/10.1126/science.1259114.

Brown, Chris A., Andrew W. Murray, and Kevin J. Verstrepen. 2010. “Rapid Expansion and Functional Divergence of Subtelomeric Gene Families in Yeasts.” Current Biology 20 (10). Elsevier Ltd: 895–903. https://doi.org/10.1016/j.cub.2010.04.027.

Chuong, Edward B, Nels C Elde, and Cédric Feschotte. 2016. “Regulatory Activities of Transposable Elements: From Conflicts to Benefits.” Nature Reviews. Genetics 18 (2): 71–86. https://doi.org/10.1038/nrg.2016.139.

Coleman-Derr, Devin, and Daniel Zilberman. 2012. “Deposition of Histone Variant H2A.Z within Gene Bodies Regulates Responsive Genes.” PLoS Genetics 8 (10): e1002988. https://doi.org/10.1371/journal.pgen.1002988.

Collier, Sarah M., Louis-Philippe Hamel, and Peter Moffett. 2011. “Cell Death Mediated by the N-Terminal Domains of a Unique and Highly Conserved Class of NB-LRR Protein.” Molecular Plant-Microbe Interactions 24 (8): 918–31. https://doi.org/10.1094/MPMI-03-11-0050.

Colome-Tatche, M., S. Cortijo, R. Wardenaar, L. Morgado, B. Lahouze, A. Sarazin, M. Etcheverry, et al. 2012. “Features of the Arabidopsis Recombination Landscape Resulting from the Combined Loss of Sequence Variation and DNA Methylation.” Proceedings of the National Academy of Sciences 109 (40): 16240–45. https://doi.org/10.1073/pnas.1212955109.

Colomé-Tatché, Maria, Sandra Cortijo, René Wardenaar, Lionel Morgado, Benoit Lahouze, Alexis Sarazin, Mathilde Etcheverry, et al. 2012. “Features of the Arabidopsis Recombination Landscape Resulting from the Combined Loss of Sequence Variation and DNA Methylation.” Proceedings of the National Academy of Sciences 109 (40): 16240–45. https://doi.org/10.1073/pnas.1212955109.

Cortijo, Sandra, Rene Wardenaar, Maria Colome, Arthur Gilly, Mathilde Etcheverry, Karine Labadie, and Fréderic Hospital. 2014. “Mapping the Epigenetic Basis of Complex Traits.” Science (New York, N.Y.) 343 (6175): 1145–48. https://doi.org/10.1038/nrg2719.

Denver, D. R., P. C. Dolan, L. J. Wilhelm, W. Sung, J. I. Lucas-Lledo, D. K. Howe, S. C. Lewis, et al. 2009. “A Genome-Wide View of Caenorhabditis Elegans Base-Substitution Mutation Processes.” Proceedings of the National Academy of Sciences 106 (38): 16310–14. https://doi.org/10.1073/pnas.0904895106.

Dobin, Alexander, Carrie A Davis, Felix Schlesinger, Jorg Drenkow, Chris Zaleski, Sonali Jha, Philippe Batut, Mark Chaisson, and Thomas R Gingeras. 2013. “STAR: Ultrafast Universal RNA-Seq Aligner.” Bioinformatics 29 (1). Oxford University Press: 15–21.

Doseff, A, R Martienssen, and V Sundaresan. 1990. “Somatic Excision of the Mu1 Transposableelement of Maize.” Nucleic Acids Research Acids Res 19 (3): 579–84.

Fu, Yu, Akira Kawabe, Mathilde Etcheverry, Tasuku Ito, Atsushi Toyoda, Asao Fujiyama, Vincent Colot, Yoshiaki Tarutani, and Tetsuji Kakutani. 2013. “Mobilization of a Plant Transposon by Expression of the Transposon-Encoded Anti-Silencing Factor.” EMBO Journal 32 (17): 2407–17. https://doi.org/10.1038/emboj.2013.169.

Gilly, Arthur, Mathilde Etcheverry, Mohammed-Amin Amin Madoui, Julie Guy, Leandro Quadrana, Adriana Alberti, Antoine Martin, et al. 2014. “TE-Tracker: Systematic Identification of Transposition Events through Whole-Genome Resequencing.” BMC Bioinformatics 15 (1). BioMed Central Ltd: 377. https://doi.org/10.1186/s12859-014-0377-z.

Gu, Muxin, Yanin Naiyachit, Thomas J. Wood, and Catherine B. Millar. 2015. “H2A.Z Marks Antisense Promoters and Has Positive Effects on Antisense Transcript Levels in Budding Yeast.” BMC Genomics 16 (1): 1–11. https://doi.org/10.1186/s12864-015-1247-4.

Henikoff, Steven, and Mitchell Mitchell Smith. 2015. “Histone Variants and Epigenetics.” Cold Spring Harbor Perspectives in Biology 7 (1). https://doi.org/10.1101/cshperspect.a019364.

Huang, Cheng Ran Lisa, Kathleen H. Burns, and Jef D. Boeke. 2012. “Active Transposition in Genomes.” Annual Review of Genetics 46 (1): 651–75. https://doi.org/10.1146/annurev-genet-110711-155616.

Ito, Hidetaka, Hervé Herve Gaubert, Etienne Bucher, Marie Mirouze, Isabelle Vaillant, Jerzy Paszkowski, Hervé Herve Gaubert, et al. 2011. “An SiRNA Pathway Prevents Transgenerational Retrotransposition in Plants Subjected to Stress.” Nature 472 (7341). Nature Publishing Group, a division of Macmillan Publishers Limited. All Rights Reserved.: 115–19. https://doi.org/10.1038/nature09861.

Ito, Tasuku, Yoshiaki Tarutani, Taiko Kim To, Mohamed Kassam, Evelyne Duvernois-Berthet, Sandra Cortijo, Kazuya Takashima, et al. 2015. “Genome-Wide Negative Feedback Drives Transgenerational DNA Methylation Dynamics in Arabidopsis.” PLoS Genetics 11 (4): e1005154. https://doi.org/10.1371/journal.pgen.1005154.

Johannes, Frank, Emmanuelle Porcher, Felipe K Teixeira, Vera Saliba-colombani, Juliette Albuisson, Fabiana Heredia, Vincent Colot, et al. 2009. “Assessing the Impact of Transgenerational Epigenetic Variation on Complex Traits.” Plos Genetics 5 (6): e1000530. https://doi.org/10.1371/journal.pgen.1000530.

Kawakatsu, Taiji, Shao shan Carol Huang, Florian Jupe, Eriko Sasaki, Robert J J. Schmitz, Mark A A. Urich, Rosa Castanon, et al. 2016. “Epigenomic Diversity in a Global Collection of Arabidopsis Thaliana Accessions.” Cell 166 (2): 492–506. https://doi.org/10.1016/j.cell.2016.06.044.

Keightley, Peter D, Urmi Trivedi, Marian Thomson, Fiona Oliver, Sujai Kumar, and Mark L Blaxter. 2009. “Analysis of the Genome Sequences of Three Drosophila Melanogaster Spontaneous Mutation Accumulation Lines,” 1195–1201. https://doi.org/10.1101/gr.091231.109.more.

Langmead, Ben, and Steven L Salzberg. 2012. “Fast Gapped-Read Alignment with Bowtie 2.” Nature Methods 9 (4): 357–59. https://doi.org/10.1038/nmeth.1923.

Li, Chenlong, Chen Chen, Lei Gao, Songguang Yang, Vi Nguyen, Xuejiang Shi, Katherine Siminovitch, et al. 2015. “The Arabidopsis SWI2/SNF2 Chromatin Remodeler BRAHMA Regulates Polycomb Function during Vegetative Development and Directly Activates the Flowering Repressor Gene SVP.” PLoS Genetics 11 (1): 1–25. https://doi.org/10.1371/journal.pgen.1004944.

Li, Guang, Shujing Liu, Jiawei Wang, Jianfeng He, Hai Huang, Yijing Zhang, and Lin Xu. 2014. “ISWI Proteins Participate in the Genome-Wide Nucleosome Distribution in Arabidopsis.” Plant Journal 78 (4): 706–14. https://doi.org/10.1111/tpj.12499.

Li, Heng. 2018. “Minimap2: Pairwise Alignment for Nucleotide Sequences.” Bioinformatics 34 (18): 3094–3100. https://doi.org/10.1093/bioinformatics/bty191.

Li, Heng, Bob Handsaker, Alec Wysoker, Tim Fennell, Jue Ruan, Nils Homer, Gabor Marth, Goncalo Abecasis, and Richard Durbin. 2009. “The Sequence Alignment/Map Format and SAMtools.” Bioinformatics 25 (16). Oxford University Press: 2078–79.

Lin, I Winnie, Davide Sosso, Li-Qing Chen, Klaus Gase, Sang-Gyu Kim, Danny Kessler, Peter M Klinkenberg, et al. 2014. “Nectar Secretion Requires Sucrose Phosphate Synthases and the Sugar Transporter SWEET9.” Nature 508 (March). Nature Publishing Group, a division of Macmillan Publishers Limited. All Rights Reserved.: 546. http://dx.doi.org/10.1038/nature13082.

Lippman, Zachary, Anne-Valérie Gendrel, Michael Black, Matthew W Vaughn, Neilay Dedhia, W Richard McCombie, Kimberly Lavine, et al. 2004. “Role of Transposable Elements in Heterochromatin and Epigenetic Control.” Nature 430 (6998): 471–76. https://doi.org/10.1038/nature02651.

Lisch, Damon. 2013. “How Important Are Transposons for Plant Evolution?” Nature Reviews Genetics 14 (1): 49–61. https://doi.org/10.1038/nrg3374.

Lloyd, John P., Alexander E. Seddon, Gaurav D. Moghe, Matthew C. Simenc, and Shin-Han Shiu. 2015. “Characteristics of Plant Essential Genes Allow for Within- and between-Species Prediction of Lethal Mutant Phenotypes.” The Plant Cell 27 (8): 2133–47. https://doi.org/10.1105/tpc.15.00051.

Love, Michael I, Wolfgang Huber, and Simon Anders. 2014. “Moderated Estimation of Fold Change and Dispersion for RNA-Seq Data with DESeq2.” Genome Biology 15 (12). BioMed Central: 550.

Lyons, David B., and Daniel Zilberman. 2017. “DDM1 and Lsh Remodelers Allow Methylation of DNA Wrapped in Nucleosomes.” ELife 6: 1–20. https://doi.org/10.7554/eLife.30674.

Makova, Kateryna D., and Ross C. Hardison. 2015. “The Effects of Chromatin Organization on Variation in Mutation Rates in the Genome.” Nature Reviews Genetics 16 (4). Nature Publishing Group: 213–23. https://doi.org/10.1038/nrg3890.

March-Díaz, Rosana, Mario García-Domínguez, Jorge Lozano-Juste, José León, Francisco J. Florencio, and José C. Reyes. 2008. “Histone H2A.Z and Homologues of Components of the SWR1 Complex Are Required to Control Immunity in Arabidopsis.” Plant Journal 53 (3): 475–87. https://doi.org/10.1111/j.1365-313X.2007.03361.x.

Marí-Ordóñez, Arturo, Antonin Marchais, Mathilde Etcheverry, Antoine Martin, Vincent Colot, and Olivier Voinnet. 2013. “Reconstructing de Novo Silencing of an Active Plant Retrotransposon.” Nature Genetics 45 (9): 1029–39. https://doi.org/10.1038/ng.2703.

Masson, Patrick, Richard Surosky, Jeffrey A Kingsbury, and Nina V Fedoroff. 1987. “Genetic and Molecular Analysis of the Spm-Dependent a- M2 Alleles of the Maize a Locus.” Genetics 177: 117–37.

Meneghini, Marc D., Michelle Wu, and Hiten D. Madhani. 2003. “Conserved Histone Variant H2A.Z Protects Euchromatin from the Ectopic Spread of Silent Heterochromatin.” Cell 112 (5): 725–36. https://doi.org/10.1016/S0092-8674(03)00123-5.

Mirouze, Marie, Jon Reinders, Etienne Bucher, Taisuke Nishimura, Korbinian Schneeberger, Stephan Ossowski, Jun Cao, Detlef Weigel, Jerzy Paszkowski, and Olivier Mathieu. 2009. “Selective Epigenetic Control of Retrotransposition in Arabidopsis.” Nature 461 (7262). Nature Publishing Group: 427–30. https://doi.org/10.1038/nature08328.

Miyao, Akio, Katsuyuki Tanaka, Kazumasa Murata, Hiromichi Sawaki, Shin Takeda, Kiyomi Abe, Yoriko Shinozuka, Katsura Onosato, and Hirohiko Hirochika. 2003. “Target Site Specificity of the Tos17 Retrotransposon Shows a Preference for Insertion Withn Genes and against Insertion in Retrotransposon-Rich Regions of the Genome.” The Plant Cell 15 (August): 1771–80. https://doi.org/10.1105/tpc.012559.ements.

Mularoni, Loris, Yulian Zhou, Tyson Bowen, Sunil Gangadharan, Sarah J. Wheelan, and Jef D. Boeke. 2012. “Retrotransposon Ty1 Integration Targets Specifically Positioned Asymmetric Nucleosomal DNA Segments in TRNA Hotspots.” Genome Research 22 (4): 693–703. https://doi.org/10.1101/gr.129460.111.

Murray, G, and William F Thompson. 1980. “Rapid Isolation of High Molecular Weight Plant DNA.” Nucleic Acids Research 8 (19). Oxford University Press: 4321–26.

Ossowski, Stephan, Korbinian Schneeberger, José Ignacio Lucas-Lledó, Norman Warthmann, Richard M Clark, Ruth G Shaw, Detlef Weigel, and Michael Lynch. 2010. “The Rate and Molecular Spectrum of Spontaneous Mutations in Arabidopsis Thaliana.” Science 327 (5961): 92–94. https://doi.org/10.1126/science.1180677.

Pietzenuk, Björn, Catarine Markus, Hervï⸮½ Gaubert, Navratan Bagwan, Aldo Merotto, Etienne Bucher, and Ales Pecinka. 2016. “Recurrent Evolution of Heat-Responsiveness in Brassicaceae COPIA Elements.” Genome Biology 17 (1). Genome Biology: 1–15. https://doi.org/10.1186/s13059-016-1072-3.

Quadrana, Leandro, Amanda Bortolini Silveira, George F. Mayhew, Chantal LeBlanc, Robert A. Martienssen, Jeffrey A. Jeddeloh, and Vincent Colot. 2016. “The Arabidopsis Thaliana Mobilome and Its Impact at the Species Level.” ELife 5 (JUN2016): 1–25. https://doi.org/10.7554/eLife.15716.

Sadeghi, Laia, Carolina Bonilla, Annelie Strålfors, Karl Ekwall, and J. Peter Svensson. 2011. “Podbat: A Novel Genomic Tool Reveals Swr1-Independent H2A.Z Incorporation at Gene Coding Sequences through Epigenetic Meta-Analysis.” PLoS Computational Biology 7 (8). https://doi.org/10.1371/journal.pcbi.1002163.

Sequeira-Mendes, J., I. Araguez, R. Peiro, R. Mendez-Giraldez, X. Zhang, S. E. Jacobsen, U. Bastolla, and C. Gutierrez. 2014. “The Functional Topography of the Arabidopsis Genome Is Organized in a Reduced Number of Linear Motifs of Chromatin States.” The Plant Cell 26 (6): 2351–66. https://doi.org/10.1105/tpc.114.124578.

Slotkin, R Keith, and Robert Martienssen. 2007. “Transposable Elements and the Epigenetic Regulation of the Genome.” Nature Reviews. Genetics 8 (4): 272–85. https://doi.org/10.1038/nrg2072.

Soppe, Wim J.J., Zuzana Jasencakova, Andreas Houben, Tetsuji Kakutani, Armin Meister, Michael S. Huang, Steven E. Jacobsen, Ingo Schubert, and Paul F. Fransz. 2002. “DNA Methylation Controls Histone H3 Lysine 9 Methylation and Heterochromatin Assembly in Arabidopsis.” EMBO Journal 21 (23): 6549–59. https://doi.org/10.1093/emboj/cdf657.

Sullivan, Alessandra M., Andrej A. Arsovski, Janne Lempe, Kerry L. Bubb, Matthew T. Weirauch, Peter J. Sabo, Richard Sandstrom, et al. 2014. “Mapping and Dynamics of Regulatory DNA and Transcription Factor Networks in A. Thaliana.” Cell Reports 8 (6): 2015–30. https://doi.org/10.1016/j.celrep.2014.08.019.

Sultana, Tania, Alessia Zamborlini, Gael Cristofari, and Pascale Lesage. 2017. “Integration Site Selection by Retroviruses and Transposable Elements in Eukaryotes.” Nature Reviews. Genetics 18 (5). Nature Publishing Group: 292–308. https://doi.org/10.1038/nrg.2017.7.

Suto, Robert K., Michael J. Clarkson, David J. Tremethick, and Karolin Luger. 2000. “Crystal Structure of a Nucleosome Core Particle Containing the Variant Histone H2A.Z.” Nature Structural Biology 7 (12): 1121–24. https://doi.org/10.1038/81971.

Teixeira, Felipe Karam, Martyna Okuniewska, Colin D. Malone, Rémi Xavier Coux, Donald C. Rio, and Ruth Lehmann. 2017. “PiRNA-Mediated Regulation of Transposon Alternative Splicing in the Soma and Germ Line.” Nature 552 (7684). Nature Publishing Group: 268–72. https://doi.org/10.1038/nature25018.

Zahraeifard, Sara, Maryam Foroozani, Aliasghar Sepehri, Dong-Ha Oh, Guannan Wang, Venkata Mangu, Bin Chen, Niranjan Baisakh, Maheshi Dassanayake, and Aaron P Smith. 2018. “Rice H2A.Z Negatively Regulates Genes Responsive to Nutrient Starvation but Promotes Expression of Key Housekeeping Genes.” J Exp Bot, no. June. https://doi.org/10.1093/jxb/ery244.

Zerbino, Daniel R., and Ewan Birney. 2008. “Velvet: Algorithms for de Novo Short Read Assembly Using de Bruijn Graphs.” Genome Research 18 (5): 821–29. https://doi.org/10.1101/gr.074492.107.

Zhu, Y. O., M. L. Siegal, D. W. Hall, and D. A. Petrov. 2014. “Precise Estimates of Mutation Rate and Spectrum in Yeast.” Proceedings of the National Academy of Sciences 111 (22): E2310–18. https://doi.org/10.1073/pnas.1323011111.

Zilberman, Daniel, Devin Coleman-Derr, Tracy Ballinger, and Steven Henikoff. 2008. “Histone H2A.Z and DNA Methylation Are Mutually Antagonistic Chromatin Marks.” Nature 456 (7218): 125–29. https://doi.org/10.1038/nature07324.

Zlatanova, Jordanka, and Amit Thakar. 2008. “H2A.Z: View from the Top.” Structure 16 (2): 166–79. https://doi.org/10.1016/j.str.2007.12.008.

